# Unravelling the plausible metal-dependent catalytic mechanism of Inositol monophosphatase ortholog from *Pseudomonas aeruginosa* through the lenses of macromolecular crystallography and enzyme kinetics

**DOI:** 10.64898/2026.04.06.716684

**Authors:** Vinay Kumar Yadav, Abinash Kumar Jena, Mitali Mukerji, Amit Mishra, Sudipta Bhattacharyya

## Abstract

The inositol monophosphatase (IMPase) orthologue is pivotal for virulence, pathogenesis, and biofilm regulation, and is therefore considered a potential drug target in *Pseudomonas aeruginosa* and other bacterial pathogens. The mammalian IMPase orthologue is an established drug target for bipolar disorder. The precise catalytic mechanism in this class of enzymes remains obscure despite five to six decades of extensive efforts and detailed studies of substrate, transition-state analogue, and product-bound structures. Here, we have solved the crystal structures of the IMPase orthologue from *Pseudomonas aeruginosa* (PaIMPase), capturing pre- and post-catalytic snapshots of metal-substrate- and metal-product-mimic-bound states. Moreover, we solved the metal-substrate transition-state-analogue-bound crystal structure of the enzyme. Critical evaluation of these high-resolution crystal structures of PaIMPase complexed with substrate, transition-state analogue, and product mimic (myo-inositol and phosphate) supports three Mg^2+^-dependent catalytic mechanisms of PaIMPase. The structural snapshots indicate that, at the enzyme active site, a metal (Mg^2+^)-coordinating water molecule, activated by two bound Mg^2+^ ions and the active-site-proximal Threonine/Aspartate dyad, attacks the central phosphorus atom of the bound substrate, leading to formation of a trigonal bipyramidal transition state. Following that, the immediate breakdown of the P-O bond results in the formation of inositolate and phosphate ions. The second water molecule, activated by another Mg^2+^ dyad, facilitates the departure of myo-inositol and phosphate from the active site. The detailed mechanistic insights gained from this work may offer a foundation for the rational design of precise inhibitors against PaIMPase.

## 1. Introduction

The inositol monophosphatases (IMPase) play vital roles in bacteria, archaea and eukaryotic organisms, including humans. It is an Mg^2+^-activated and Li^+^-sensitive enzyme which often demonstrates substrate promiscuity depending on the host organism. IMPase orthologs have quite different functionalities in bacteria, including those that survive at high temperatures (Involved in osmolyte production). In lower-order organisms, IMPase orthologues are involved in pathways that regulate virulence factors, including biofilm formation in *Burkholderia* cenocepacia (*B. cenocepacia*) and *Staphylococcus aureus* (*S. aureus)*, exopolysaccharide biosynthesis in Rhizobium *leguminosarum* and *S. aureus*, etc. (Rosales-Reyes *et al*., 2012; Janczarek & Skorupska, 2001; Boles *et al*., 2010; Dutta *et al*., 2014). In mycobacteria, these enzymes synthesise mycothiol, an alternate of glutathione as a defence strategy of the pathogen against reactive oxygen and nitrogen species (Movahedzadeh *et al*., 2010). The role of this enzyme is extensively explored in *E. coli,* suggesting its involvement in cold sensitivity and acting as an extragenic suppressor of the temperature-sensitive phenotypes. Additionally, as a part of the transcription antitermination complex, it interacts with NusA and subsequently regulates 30S ribosome biogenesis, rRNA folding and maturation (Wang *et al*., 2007; Dudenhoeffer *et al*., 2019; Singh *et al*., 2016; Huang *et al*., 2019). IMPase ortholog, SuhB in *Pseudomonas aeruginosa* (*P. aeruginosa*) has a very diverse functional implications, including regulation of expression of Type III secretion system (T3SS) proteins, swimming motility of the bacteria, host cell cytotoxicity during acute infection, bacterial biofilm formation during chronic infections and the regulation of antimicrobial drug resistance of the pathogen (Li *et al*., 2013, 2017; Shi *et al*., 2015). The functional implications of the IMPases are not limited to the lower-order organisms; instead, it regulates the normal physiology of the human central nervous system. Myo-inositol (MI), present in relatively high concentrations in the brain, is a precursor for various cell-signalling molecules, including phosphatidylinositol and inositol phosphates, which play crucial roles in essential cellular processes. Strict regulation of MI and its phosphate levels in the brain is crucial, as any alteration may lead to a severe brain disorder. Brain cells have well-organised regulatory networks for the de novo synthesis, transport, and recycling of MI and its derivatives. As the blood-brain barrier permeability of MI is very poor, the main source of MI in the brain is its synthesis through the de novo pathway. In MI biosynthesis pathways, the prominent role of IMPase is to convert myo-inositol-1-phosphate (IPD) into MI and inorganic PO_4_^3-^. The MI synthesis pathway is hyperactive in bipolar disorder, which poses significant global impacts on the healthcare industry (Raghu *et al*., 2019; Spector, 1988; Shaltiel *et al*., 2001). To date, the 3D structures of the IMPases from different organisms have been solved and studied to understand the organism-specific structural and functional aspects of these enzymes. X-ray diffraction crystallography, NMR and single particle cryo-electron microscopy have been extensively applied to solve 3D structures of IMPases in their apo as well as different metal ions, with their substrates and product-bound forms. Structurally, IMPases mostly form a homodimer composed of ∼30 kDa subunits. Each protomer subunit is built from alternating layers of α-helices and β-sheets, arranged into a five-layered αβαβα sandwich. This conserved αβαβα core contains 9 α-helices and 13 β-strands, a feature also strongly preserved in fructose 1,6-bisphosphatases (FBPase) and inositol polyphosphate- 1-phosphatases (IPPase). The IMPase enzyme’s active site lies within a hydrophilic cavity bordered by α2, β3, α8, and residues 90–95. At the cavity entrance, a β-hairpin structure (β1: Ile33–Lys36; β2: Asp41–Thr44) provides additional architectural definition. The structural studies collectively reveal that there are three metal-binding sites in each active site of IMPase monomeric structure. However, each metal binding sites have different binding affinity and plays a distinct role individually or in combination (York *et al*., 1995; Ke *et al*., 1989; Bone *et al*., 1994*a*; Li *et al*., 2010). The crystal structure of human IMPase (PDB codes 1IMA and 1IMB) indicates the presence of a substrate complexed with a single metal ion (Gd^3+^), corroborating the existence of a higher-affinity metal-binding site. Moreover, the presence of three Mn^2+^ in apo human IMPase structure (PDB code: 1IMC), three Ca^2+^ bound D-Ins (1)P (PDB ID:1AWB) and two Ca^2+^ in archaeal IMPase (PDB code: 1G0H) reveals the 3D disposition of each metal and also suggests that two metal ions are sufficient for enzyme catalysis (Pollack *et al*., 1994; Bone *et al*., 1994; Johnson *et al*., 2001; Gill *et al*., 2005; Arai *et al*., 2007). Furthermore, a few in silico-based studies also concluded that the presence of two metal ions is energetically more favourable with one water molecule bound by the second metal, assisting in the protonation of leaving products. Conversely, the requirement of a third metal ion is necessary for water nucleophile activation and initiation of dephosphorylation, also concluded by another in silico-based study (Wang & Hirao, 2013; Lu *et al*., 2012).

The structural study of the human IMPase broadly focuses on the coordination geometry and coordination numbers of the bound metals at the active sites. Within the IMPase active site, Mg²⁺ binding site 1 (Mg²⁺-1) is formed by the carboxylates of Glu70 and Asp90, the carbonyl group of Ile92, and three water molecules. Mg²⁺ binding site 2 (Mg²⁺-2) involves the carboxylates of Asp90, Asp93, Asp220, and three water molecules, one of which is shared with Mg²⁺-1. Mg²⁺ binding site 3 (Mg²⁺-3) consists of a single carboxylate from Glu70 and five water molecules, one of which is also shared with Mg²⁺-1. Because Mg²⁺-3 interacts only weakly with IMPase, it is occupied only at higher Mg²⁺ concentrations. Notably, Mg²⁺-1 and Mg²⁺-2 correspond to the binding sites of the two Mn²⁺ ions and function analogously to the two-metal-ion mechanism. The role of the highly conserved Thr/Asp dyad, along with the third metal in water nucleophile activation, was also suggested by an in-silico study (Lu *et al*., 2012*b*). Recent data on metal ion affinity for optimal catalysis in archaeal IMPase, obtained through isothermal titration calorimetry and further supported by the active site mobile loop mutants, strongly support a three-metal-ion-assisted mechanism. Consequently, the previously proposed two-metal-ion mechanism for FBPase/IMPase has been revised to a three-metal-ion model. Moreover, the role of the α4 helix and its preceding loop is extensively explained by Bhattacharya et al., suggesting that the length of this helix directly determines the substrate specificity among this class of enzymes. The mechanism of Li^+^ ion-based inhibition is also extensively studied, which suggests that Li^+^ occupy the second metal site, subsequently causing product trapping due to the unavailability of protonating water enclosures by the tetrahedral geometry of Li^+^ as compared to the octahedral geometry of Mg^2+^. Additionally, Li^+^ also interferes with the positioning of the active site mobile loop, which may lead to the unavailability of activated water nucleophile to initiate the enzymatic catalysis of the bound phosphate ester. Despite all these aforementioned atomic-level observations, until now, there is no availability of the transition-state analogue-bound 3D structural coordinates of this class of enzymes, which is a major lack in this field. Apart from that, there is no availability of apo-, substrate-, transition state analogues-, and product-bound crystal structures of an IMPase from a single organism, which is indispensably needed to understand the catalytic mechanism comprehensively from the beginning of the catalytic events to the end of catalysis. This manuscript aims to conduct a comprehensive study explaining the mechanism of the enzyme catalysis of the IMPases, taking PaIMPase as a reference member of this family, using high-resolution X-ray crystallography. Therefore, we have applied the enzyme kinetics and macromolecular crystallography to bridge the research gaps mentioned earlier in this section. We have performed a basic enzyme-inhibition study and solved the crystal structures of metal-supplemented PaIMPase in complex with its substrates (2’AMP), the substrate transition state analogue (Sodium tungstate), and products (MI and PO_4_^3-^) to delineate every step of the catalytic events through the lens of structural biology and visual observation of different molecular snapshots we also analysed the PaIMPase apo (PDB ID: 8WIP) and IPD-bound (PDB ID: 8WDQ) crystal structures published elsewhere (Yadav *et al*., 2025). These findings will provide a more concrete view of catalytic events and help to design more precise and selective inhibitors of this class of enzymes, which may subsequently help to eradicate bipolar disorder and life-threatening bacterial infections.

## 2.0. Materials and Methods

### 2.1. Cloning and purification of PaIMPase

*Paimpase* gene (PA3818) was PCR amplified from the genomic DNA of *P. aeruginosa* PAO1 strain as template using Taq master mix and gene-specific forward and reverse primers flanked with *Bam*HI and *Hin*DIII restriction sites at 5’ and 3’ respectively, followed by the purification of the PCR product using PCR clean up kit as per manufacturer protocol of QIAGEN. Restriction digestion of the PCR product and pET28a vector with respective enzymes was performed, followed by purification and then ligation of insert and vector at a ratio of 5:1. The chemically competent *E. coli* DH5 α cells were transformed with ligation mixtures, and positive clones were selected on kanamycin-containing LB agar plates followed by confirmation of positive clones via colony PCR, restriction digestion and gene sequencing (Inoue *et al*., 1990). The overexpression of PaIMPase protein was attempted in *E. coli* BL21(DE3) expression strain at 16 ℃ and 37 ℃ for 4 hrs and 16 hrs respectively, with 100 µM IPTG as inducer in the presence of an appropriate amount of kanamycin. Since recombinantly overexpressed PaIMPase was fairly soluble at 37℃ with an ample amount of protein, as indicated by SDS-PAGE, we proceeded to purify the recombinant protein using large volumes of culture. Post 4.5 hrs of IPTG induction, cells were harvested at 4500 rpm for 30 minutes in a swing bucket rotor centrifuge, followed by cell lysis via sonication in buffer (10 mM Tris-base, 300 mM NaCl, 10% glycerol and 10 mM imidazole, pH 8.0) and then centrifugation at 12k/90 min/4 ℃. The cytosolic fraction was subjected to protein purification through pre-equilibrated Ni-NTA-Sepharose beads due to the presence of poly-His tag at the N-terminal. The recombinant protein was eluted using an increasing imidazole concentration in the same buffer. Presence of desired protein was confirmed by SDS-PAGE on 12% acrylamide gel. The protein purified through nickel affinity was subjected to gel filtration chromatography using AKTA start for further purification and buffer exchange. The composition of the gel filtration buffer was 10 mM Tris-base, 100 mM NaCl and 5 mM DTT, pH 8.0.

### 2.2. Biochemical characterisation of IMPase activity

To test the enzyme activity of the purified protein, first we have examined the substrate specificity against different plausible substrates including myo-D-inositol-1-monophosphate, adenosine-2′-monophosphate, p-nitrophenyl phosphate, oxidized and reduced form of nicotinamide adenine dinucleotide phosphate, fructose-1, 6-bisphosphate, adenosine-5′-phosphate, adenosine-3′, 5′-cyclic monophosphate, and adenosine 3′, 5′-bisphosphate **(Figure 1A**) in the presence of 10 mM Mg^2+^ ions and buffer containing 20 mM Tris pH 8.0 and 150 mM NaCl, followed by incubation of reaction mixture at 37 ℃ for 2 minutes. The product formation was determined by using the malachite green phosphate assay kit with slight modifications in the protocols (Feng *et al*., 2011). A similar reaction condition was applied throughout all biochemical characterisation unless otherwise specified. We have performed divalent cation specificity check of the enzyme in the presence of 10 mM of each cation separately (Given in **Figure 1B**) and 50 µM of IPD as substrate. Since we found Mg^2+^ is the most preferred cofactor, we performed optimisation of Mg^2+^ concentration for maximum activity in the presence of varying Mg^2+^ concentrations. The kinetic parameters, like K_m_ and V_max_ for the two substrates, IPD and 2′AMP, were examined in the presence of 30 mM Mg^2+^ and varying concentrations of substrates. Enzyme activity abolition at higher concentration of Mg^2+^ was also investigated, similarly with varying concentration of Mg^2+^ till 1 M. Li^+^ based IMPase activity inhibition of PaIMPase was also attempted in the presence of 30 mM of Mg^2+^ and varying concentration of Li^+^ ions with the same reaction conditions. All the graphs were plotted with Origin and GraphPad Prism.

**Figure 1:**
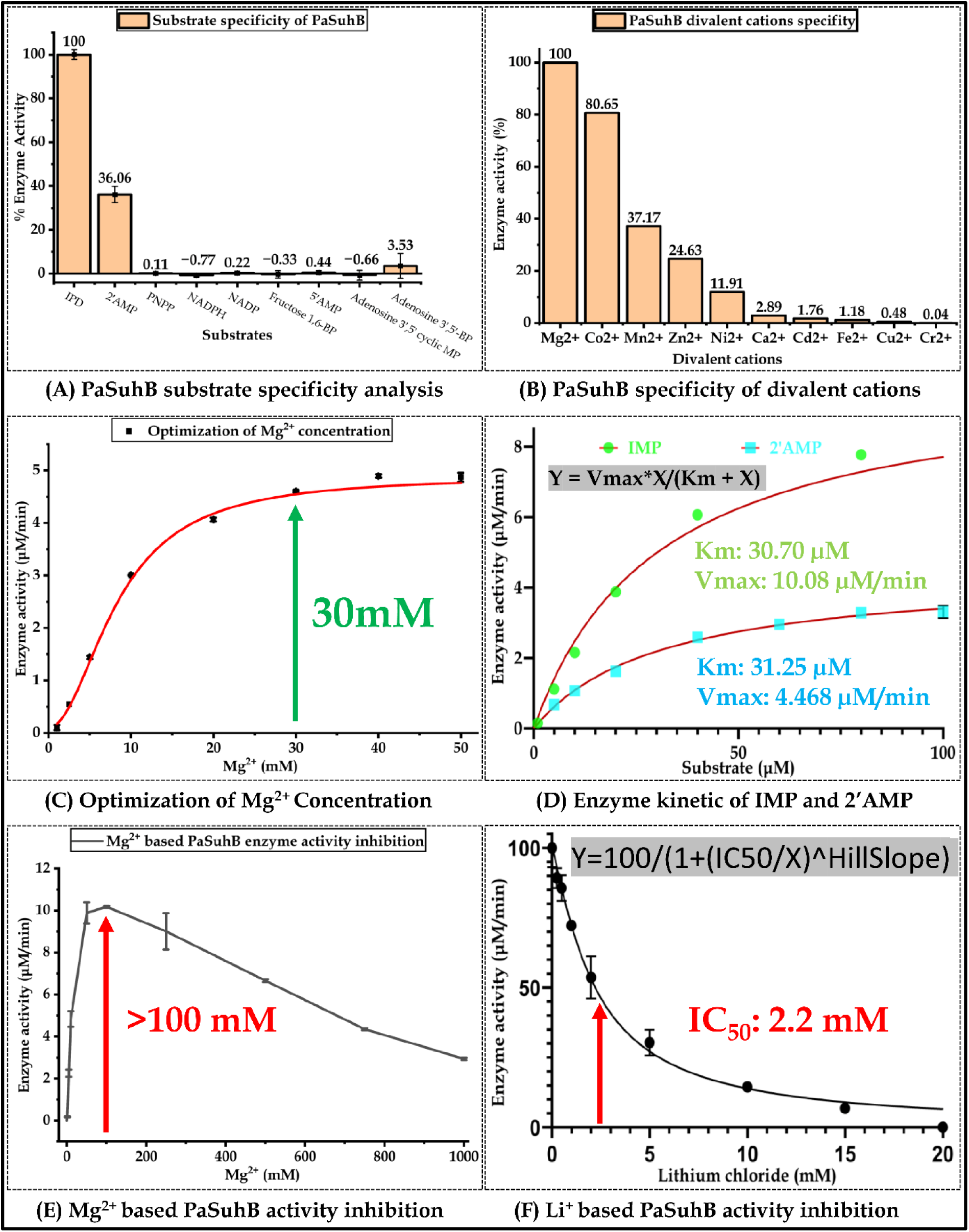
IMPase-like characteristic of PaIMPase. Checking A) Substrate specificity, B) Divalent cations-specificity, C) Optimisation of Mg^2+^ concentration for maximum enzymatic activity, D) Evaluation of kinetic parameters for two most preferred substrates IPD and 2′AMP, E) Mg^2+^ and F) Li^+^ dependent enzyme activity inhibition respectively.

### 2.3. Inhibition of PaIMPase IMPase activity with transition state analogues, sodium tungstate and sodium orthovanadate

We have used two metal-based chemical entities that structurally mimic the trigonal bipyramidal (TBP) transition state of phosphate monoester, which should hypothetically form during the hydrolysis of inositol monophosphate substrate by the enzymatic catalysis of the inositol monophosphatase enzyme. Since a good transition state analogue should closely mimic the transition state of the real substrate, it should ideally act as a competitive inhibitor with respect to the incoming substrate to inhibit its hydrolysis. Hence, to identify the best transition state analogue of PaIMPase, we have investigated the inhibition of PaIMPase enzyme activity in the presence of varying concentrations of two TBP mimicking metal based transition state analogues by varying sodium orthovanadate (up to a maximum concentration of 20 mM) and Sodium tungstate dihydrate (up to a maximum concentration of 2 mM), separately in the presence of 50 µM IPD as the substrate, 30 mM Mg^2+^ as the activating metal cofactor, in the identical reaction conditions as applied for biochemical characterization.

### 2.4. Protein crystallisation

We have attempted crystallisation of PaIMPase in the presence of its substrates (2’AMP) as well as in the presence of the metal-based transition state analogues (sodium tungstate dihydrate) and in the presence of IPD hydrolysis products [myo-inositol MI) and phosphate (PO_4_^3-^) bound forms. The crystallisation was performed in the presence of the inhibitory divalent metal ions such as Ca^2+^ for the substrates (to prevent substrate hydrolysis) and either in the presence of Mg^2+^or Ca^2+^ (whichever is compatible for crystallisation of the transition state analogue and product-bound forms). The final protein purification for crystallisation purposes was done in either buffer A (10 mM Tris-buffer, pH 8.0, 50 mM NaCl, 10 mM CaCl_2_·2H_2_O and 2 mM DTT) or buffer B (Buffer A except CaCl_2_·2H_2_O is replaced with an equal concentration of MgCl_2_·6H_2_O) with the addition of excess DTT (50 mM) during the final step of protein concentration. The apo form of PaIMPase crystals appeared in a sitting drop condition with 20 mg/ml protein in 0.1M CH3COONa · 3H2O, pH 4.5 and 5.0, with 25% PEG3350 after two days of incubation at room temperature (Yadav *et al*., 2025). Thus, to obtain the cocrystals with substrates, transition state and products, we have applied a similar strategy and almost similar crystallisation conditions. To obtain diffraction-quality single crystals with the aforementioned ligands, we have performed fine-tuning of crystallisation conditions for each ligand separately. To obtain 2’AMP-bound PaIMPase crystals, the protein was mixed with a 5X concentration of 2’AMP dissolved in the same buffer. Thick rod-like single crystals appeared in 0.1M sodium acetate trihydrate, pH 5.0, with 14 % PEG3350 post 21 days of crystal setup. To obtain the tungstate-bound PaIMPase crystals, we performed an initial crystallisation trial using 20 mg/ml of protein with a 10X concentration of sodium tungstate dihydrate dissolved in water, which resulted in the formation of microcrystals after 2 days in 0.2 M ammonium sulfate or 0.2 M lithium sulfate monohydrate, 0.1 M HEPES pH 7.5 and 25 % PEG 3350. Diffraction of these crystals reveals that they are salt crystals; hence, we tried soaking PaIMPase apo crystals (protein purified in the presence of Mg^2+^) in 0.1 M sodium acetate, pH 5.0, with 20 % PEG3350. Thus, obtained crystals were soaked in mother liquor containing an additional 100 mM MgCl_2_·6H_2_O, 20 mM sodium tungstate dihydrate and 10% glycerol as cryoprotectant for just 3.0 minutes. The crystals were soaked for a very short period of time to avoid melting of the crystals and poly-tungstate formation at acidic pH. To obtain product-bound crystals, the PaIMPase protein was purified in buffer B, followed by mixing with a 50 mM concentration of each MgCl_2_·6H_2_O, myo-inositol and sodium phosphate monobasic anhydrous. Thick rod-shaped diffraction quality crystals appeared after two days of incubation in 0.1 M sodium acetate, pH 5.0, with various PEG3350 concentrations ranging from 18-22 %.

### 2.5. Crystal diffraction, data collection and structure solution

The diffraction data collection of all the crystals was performed at the synchrotron X-ray radiation source with X-ray wavelength of 0.9732 Å (Indus 2, Beamline-21-Px, RRCAT, Indore, India). To obtain the 2’AMP-bound PaIMPase crystal structure, multiple single crystals were picked and soaked for 20 minutes in mother liquor containing 10 % glycerol, 40 mM CaCl_2_·2H_2_O and 20 mM 2’AMP. Data for the best diffracted crystal was collected for 280 frames with 1 degree oscillation width per frame, with the detector at 240 mm distance. In the context of PaIMPase tungstate-bound crystals, data collection for the best diffracted crystals from the soaking was done for 180 frames, keeping 1 degree oscillation width per frame with the detector distance at 240 mm. Since co-crystallised PaIMPase crystals with the IPD hydrolysis products, MI and PO_4_^3-^ did not show any density for these ligands at the enzyme active sites, crystals from the same crystallization drops were first soaked in mother liquor containing 200 mM CaCl_2_·2H_2_O and 10% glycerol as cryoprotectant for just 30 seconds, picked, and followed by another round of soaking in mother liquor containing 100 mM each MI and PO_4_^3-^ for ∼ 2 minutes before snap freezing. Data collection for the best diffracted crystals was done for 190 frames and 1 degree oscillation width per frame with the detector distance of 240 mm. After collecting the full-length data, the processing was done with a series of data reduction and data processing software tools. Initially, the data were reduced by indexing using XDS software (Kabsch, 2010). The reduced data were then subjected to merging and scaling by Pointless and Scala from the CCP4 package (Agirre *et al*., 2023). The content of the crystal asymmetric unit was determined by calculation of the Matthews coefficient (Matthews, 1968). Molecular replacement was done with Phaser using a single chain of PaIMPase apo structure (PDB ID 8WIP) as the search template. After obtaining the initial phase information from molecular replacement, the phase improvement was carried out through iterative cycles of model building and structure refinement using Coot (Emsley *et al*., 2010), Refmac5 or Phenix (Murshudov *et al*., 2011; Adams *et al*., 2010), respectively, until a good agreement between the experimental data and the structural model was achieved. Finally, the molecular model underwent geometrical stereochemical and other structural validation before being submitted to the Protein Data Bank (PDB; https://www.rcsb.org/). The PDB IDs for all four submitted crystals are listed in Table 1.

**Table 1:** Crystallisation parameters of 2**′**AMP, GTG and MI-PO_4_^3-^ bound PaIMPase structures.

| Parameters | PaIMPase 2'AMP complex (9WBP) | PaIMPase-Mg<sup>2+</sup>- Tungstate complex (22HF) | PaIMPase-Ca<sup>2+</sup>- MI-PO<sub>4</sub><sup>3-</sup> (9XCK) |
| --- | --- | --- | --- |
| Resolution range | 48.07 - 2.0 (2.072 - 2.0) | 39.07 - 2.55 (2.641 - 2.55) | 43.21 - 2.3 (2.382 - 2.3) |
| Space group | P 2<sub>1</sub> 2<sub>1</sub> 2<sub>1</sub> | P 2<sub>1</sub> 2<sub>1</sub> 2<sub>1</sub> | P 2<sub>1</sub> 2<sub>1</sub> 2<sub>1</sub> |
| Unit cell | 65.696 90.051 113.377 90 90 90 | 64.59 89.483 98.137 90 90 90 | 65.006 88.904 98.878 90 90 90 |
| Total reflections | 89739 (7627) | 37884 (3685) | 49891 (4977) |
| Unique reflections | 45675 (4137) | 19121 (1865) | 25793 (2549) |
| Multiplicity | 2.0 (1.8) | 2.0 (2.0) | 1.9 (2.0) |
| Completeness (%) | 96.73 (91.02) | 99.81 (99.95) | 98.68 (99.76) |
| Mean I/sigma(I) | 14.63 (2.03) | 8.99 (1.96) | 12.99 (2.99) |
| Wilson B-factor | 31.40 | 38.16 | 33.82 |
| R-merge | 0.02929 (0.2812) | 0.07658 (0.4083) | 0.04739 (0.2929) |
| R-meas | 0.04142 (0.3977) | 0.1083 (0.5775) | 0.06703 (0.4142) |
| R-pim | 0.02929 (0.2812) | 0.07658 (0.4083) | 0.04739 (0.2929) |
| CC1/2 | 0.999 (0.867) | 0.993 (0.744) | 0.997 (0.833) |
| CC* | 1 (0.964) | 0.998 (0.924) | 0.999 (0.953) |
| Reflections used in refinement | 44662 (4134) | 19103 (1864) | 25773 (2547) |
| Reflections used for R-free | 2248 (224) | 964 (114) | 1208 (117) |
| R-work | 0.2033 (0.3030) | 0.2263 (0.3132) | 0.1847 (0.2180) |
| R-free | 0.2541 (0.3696) | 0.2849 (0.4312) | 0.2413 (0.2475) |
| CC (work) | 0.952 (0.855) | 0.944 (0.795) | 0.960 (0.875) |
| CC (free) | 0.953 (0.882) | 0.904 (0.726) | 0.960 (0.965) |
| Number of non-hydrogen atoms | 4802 | 4333 | 4495 |
| Macromolecules | 4146 | 4124 | 4163 |
| Ligands | 56 | 39 | 45 |
| Solvent | 600 | 170 | 287 |
| Protein residues | 539 | 540 | 542 |
| RMS (bonds) | 0.007 | 0.003 | 0.004 |
| RMS (angles) | 1.24 | 0.74 | 0.70 |
| Ramachandran favoured (%) | 97.37 | 97.76 | 97.96 |
| Ramachandran allowed (%) | 2.63 | 2.24 | 1.86 |
| Ramachandran outliers (%) | 0.00 | 0.00 | 0.19 |
| Rotamer outliers (%) | 0.48 | 2.88 | 1.66 |
| Clash score | 4.66 | 11.86 | 5.01 |
| Average B-factor | 41.41 | 47.01 | 42.32 |
| Macromolecules | 40.13 | 47.18 | 42.07 |
| Ligands | 41.01 | 50.17 | 57.89 |
| Solvent | 50.25 | 42.25 | 43.46 |
| Number of TLS groups | 1 | 2 | 2 |

## 3. Results

### 3.1. PaIMPase exerts classical IMPase-like enzymatic characteristics

The inositol monophosphatase class of enzymes, irrespective of the organism of origin, exhibits classical catalytic behaviours, including a conserved active site pocket, sensitivity to magnesium ions, Li^+^-based inhibition, and promiscuous substrate specificity. The biochemical behaviours of IMPases in particular organisms determine their functional implications in regulating survival, pathogenesis and virulence factors. In this context, to probe the plausible enzyme-based functional aspects of PaIMPase, which is also named as SuhB (PaIMPase), we have performed multiple biochemical experiments using a malachite green phosphate assay kit to measure liberated free phosphate during catalysis. The preliminary substrate screening results (**Figure 1A**) reveal that PaIMPase is enzymatically active against IPD and 2′AMP; among these two, IPD has emerged as the most preferred substrate, as evidenced by the highest catalytic efficiency in the enzymatic assay. The divalent metal ion specificity was investigated with 10 mM concentration of probable ions, including Mg^2+^, Co^2+^, Mn^2+^, Zn^2+^, Ni^2+^, Ca^2+^, Cd^2+,^ Fe^2+^, Cu^2+^ and Cr^2+,^ in the presence of IPD as substrate (**Figure 1B**). Consistent with the behaviour of classical IMPases, PaIMPase exhibited the highest activity in the presence of Mg^2+^, suggesting that Mg^2+^ serves as a preferred divalent cation cofactor. Nonetheless, the enzyme retains catalytic activity in the presence of several other divalent cations, including Co^2+^, Mn^2+^, Zn^2+^ and to a minor extent Ni^2+^. Moreover, the optimisation of Mg^2+^ concentration (**Figure 1C**) revealed that PaIMPase exhibits maximal enzyme activity at 30 mM. The apparent affinity constant (K_a_) for Mg^2+^ was estimated to be approximately 8.38 mM. Furthermore, using optimal concentration of cofactor Mg^2+^ (30 mM), we have analysed the kinetic parameters of the two most preferred substrates, IPD and 2′AMP, in identical reaction conditions (**Figure 1D**). The results indicate that the kinetic parameters like K_m_ and V_max_ for IPD are 30.70 µM and 10.08 µM/min, while for 2′AMP are 31.25 and 4.468 µM/min, respectively. This observation signifies that PaIMPase has equal affinities for both substrates, while its catalytic efficiency is higher with IPD.

One of the most crucial observations with classical IMPase is that it exhibits reduced enzyme activity in the presence of an excess amount of Mg^2+^ ions. To investigate these characteristics in PaIMPase, we have performed Mg^2+^ concentration-dependent enzyme activity of PaIMPase in identical reaction conditions using IPD as substrate. Interestingly, our results (**Figure 1E**) indicate that, like other classical IMPases, PaIMPase also exhibited a similar characteristic, which is reduced enzyme activity at Mg^2+^ concentrations >100 mM. Additionally, the Li^+^ sensitivity of PaIMPase was investigated (**Figure 1F**) using IPD as substrate and 30 mM Mg^2+^ as activator; likewise, classical IMPases, the PaIMPase also exhibited Li^+^ based inhibition with an IC_50_ value approximately 2.2 mM. Collectively, the biochemical characterisation experiments of PaIMPase unfold that it belongs to the classical Mg^2+^ activated and Li^+^ inhibited IMPase family protein with substrate preferences towards IPD and 2′AMP, while Mg^2+^ is the most preferred cofactor.

### 3.2. Intermediate state analogues: Tungstate exerts greater potential to inhibit PaIMPase activity

To understand the catalytic mechanism of the IMPases, we need the structural snapshots of the different poses of the enzymatic reaction, like in substrate-, transition state analogue- and product-bound states of the enzyme. From the initial biochemical characterisation, it was observed that two substrates, IPD and 2’AMP, are efficiently hydrolysed by PaIMPase. Before starting crystallisation trials, we performed the enzyme activity inhibition of PaIMPase with two probable transition state analogues, sodium orthovanadate and sodium tungstate. Since these two compounds in solution may adopt a very similar trigonal bipyramidal (TBP) conformation that the substrate monoester PO_4_ ^3-^ group should attain after the in-line nucleophilic attack (through S_N_2 mechanism) of the metal (Mg^2+^) activated water molecule to the central phosphorus atom of the enzyme-bound phosphate monoester substrate during the intermediate step of the catalytic reaction. We have performed the PaIMPase inhibition reaction using the substrate, IPD (50 µM), in the presence of activator metal ion Mg^2+^ (30 mM); the concentration of the enzyme was 0.25 µM, and the rest of the reaction conditions were kept similar to the conditions mentioned for biochemical characterisation. Our results (**Figure 2**) indicate that both the transition state analogues have different potentials to inhibit the PaIMPase enzyme activity. **Figure 2A** indicates the enzyme activity inhibition profile of sodium orthovanadate, while **Figure 2B** indicates the sodium tungstate-based PaIMPase inhibition profile. These results indicate that sodium tungstate is approximately sixty-five times (IC_50_: 0.123 mM) more potent inhibitor of PaIMPase enzyme activity compared to sodium orthovanadate (IC_50_: 8.014 mM). The difference in inhibition potential and IC_50_ values indicates that in solution, sodium tungstate may interact at the PaIMPase active site more efficiently; therefore, to get the crystal structure of PaIMPase-transition-state-analogue binary complex, we have performed further PaIMPase crystallisation trials in the presence of sodium tungstate and the activator metal ion Mg^2+^.

**Figure 2:**
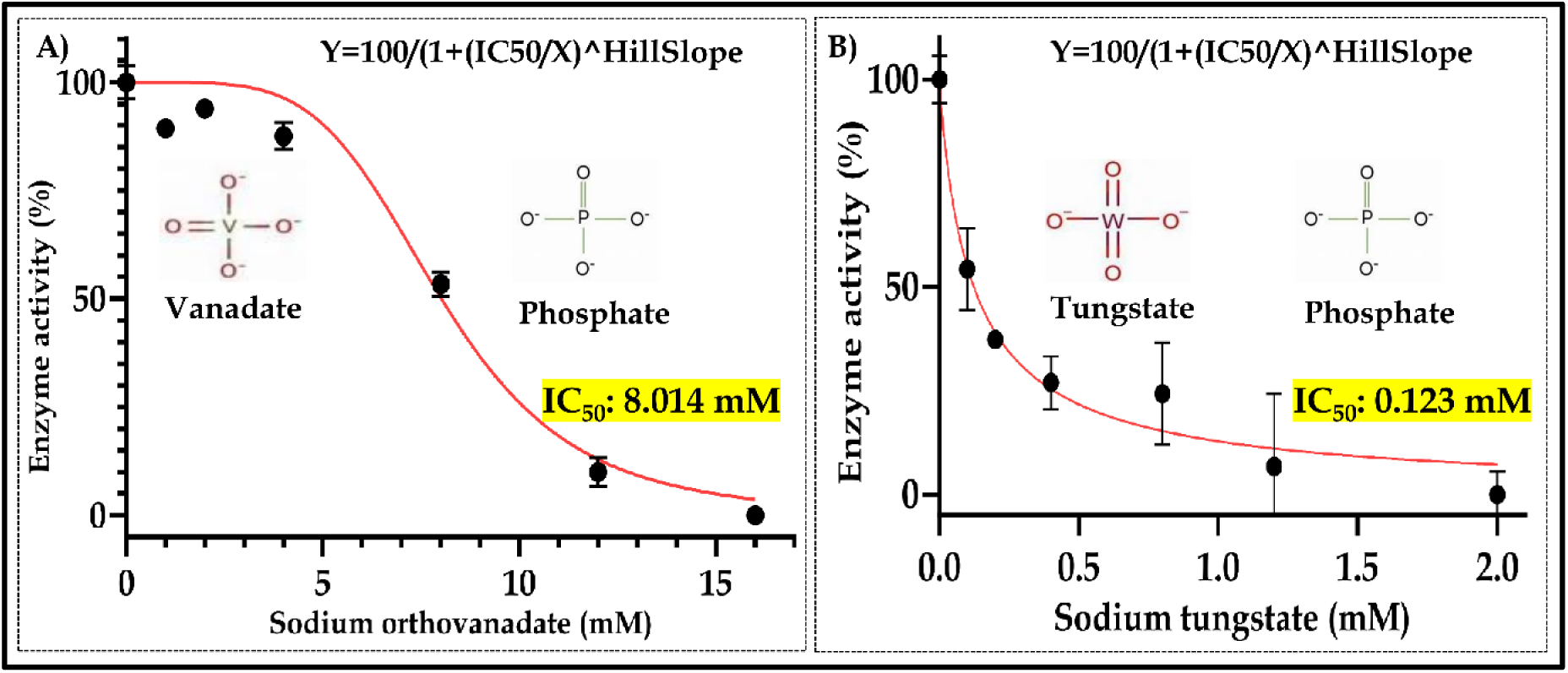
Enzyme activity inhibition of PaIMPase with transition state analogues, A) Sodium orthovanadate and B) Sodium tungstate dihydrate. The black dotted graph with error bar indicates the mean of the actual data, and the red line graph indicates the Hill fitting of the respective graphs.

### 3.3. Crystal structure of substrate (2′AMP) bound PaIMPase

The single diffraction quality crystal of PaIMPase-Ca^2+^-2′AMP complex was picked from the crystallisation condition containing 0.1 M sodium acetate trihydrate, pH 5.0, with 14 % (w/v) PEG 3350 (**Supplementary Figure S1A1**). The picked single crystal was further soaked in the same mother liquor supplemented with 40 mM CaCl_2_·2H_2_O, 20 mM 2′AMP and 10 % (V/V) glycerol as cryoprotectant. A typical diffraction frame of this crystal is shown in **Supplementary Figure S1B1**. After a full set of X-ray diffraction data collection, the data were indexed and scaled in the P2_1_2_1_2_1_ space group with unit cell dimensions: a=65.696, b=90.051, c=113.377, α=β=γ=90^ο^. The asymmetric unit of the crystal contains a single dimer, with a Matthew’s coefficient of 2.79 and corresponding estimated solvent content of 56.02 % (**Figure 3A**). The diffraction data could be rationally processed up to 2.0 Å following all the quality assurance parameters of data reduction (Table 5.1). The atomic coordinates of the structural model were refined until the R and R_free_ values were merged up to 0.2035 and 0.2541, respectively. The X-ray diffraction data collection statistics and the refined model’s quality assessment parameters fall within the expected values (**Table 1 and Supplementary Figure S2A1**). The overall dimer of PaIMPase-Ca^2+^-2′AMP complex was found to be stabilised by 18 hydrogen bonds and 21 salt bridges formed between the monomers. The interface area of the dimer is 1685.5 Å^2^ (**Supplementary Figure S2B1**). Similar to PaIMPase apo and IPD-bound structures (Yadav *et al*., 2025), the monomeric structures of each dimer of PaIMPase/Ca^2+^/2′AMP complex (**Figure 3B**) preserve its classical penta-layered sandwich architecture of alternating α and β secondary structural layers (αβαβα). This architecture is also common in other proteins of this family, including CysQ FBPase and GlpX. The constitution of the active site architecture of PaIMPase-Ca^2+^-2′AMP complex is also similar to the IPD-bound crystal structure of the protein. Unlike PaIMPase/Ca^2+^/IPD structure, both monomers of PaIMPase-Ca^2+^-2′AMP complex are very similar (RMSD: 0.093 Å) to each other **(Supplementary Figure S3**) with respect to the numbers of active site-bound metal ions and the orientation of the substrate molecule and the relative disposition of the active site mobile loop (Arg25 to Glu42), because of this inter-monomeric structural resemblance of PaIMPase-Ca^2+^-2′AMP complex, only one monomer of the complex is depicted in the **Figure 3B**. The average B-factor analysis indicates that the 2′AMP-bound crystal structure (average B-factor 41.41) is less stable and suffers from more thermal vibrations than its IPD-bound form (average B-factor 27.64), while almost equally stable to the apo structure of the protein (average B-factor 40.31) (Yadav *et al*., 2025). In the case of the 2′AMP-bound form, the active site of the protein is stabilised by four calcium ions, two molecules of 2′AMP and one molecule of glycerol. Both the active sites are occupied by one molecule of 2′AMP (**Figure 3B**). The 2Fo-Fc omit map (contoured at 1σ) of Ca^2+^ and 2′AMP, bound at the active site of PaIMPase, is represented in **Figure 3C**. Each of these bound metal ions and the substrate molecules could be refined to full (100%) occupancy. Both the Ca^2+^ ions at the active sites attain the distorted octahedral coordination geometry by interacting with the highly conserved metal binding residues residing at the active site of PaIMPase monomers, as well as the bound substrate and neighbouring water molecules. The Ca^2+^-1 ion forms six coordinate bonds, three from Asp86, Leu88, and Glu67 amino acids, one from the substrate, and two from the water molecules denoted as W1 and W3. Another Ca^2+^-2 ion also forms six bonds, three from amino acids, Asp86, Asp89 and Asp216, two from substrate and one from a water molecule denoted as W2 (**Figure 3D** and **Supplementary Table 1**). Overall, in the case of the PaIMPase-Ca^2+^-2′AMP complex, the active site catalytic cleft is formed by the amino acid residues, including Glu67, Asp86, Leu88, Asp89, Gly90, Thr91, Phe161, Arg162, Arg188, Gly190, Ala191, Ala192, Glu209, Leu212 and Asp216 (**Figure 3E-F**). The first calcium (Ca^2+^1 in purple colour 2D **Figure 3F**) interacts with Glu67, Asp86 and Leu88 and needs the O3 atom of PO_4_^3-^ moiety of 2′AMP to support its binding. The second calcium (Ca^2+^2 in purple colour 2D **Figure 3F**) interacts with three amino acid residues Asp86, Asp89 and Asp216. This calcium supports the binding of the incoming substrate 2′AMP through binding with the O2 and O3 atoms of the PO_4_^3-^ moiety of the substrate. The chain A-bound two Ca^2+^ ions have B-factors of 42.44 and 49.92 Å^2,^ which corroborates their relative stability and may reflect their relative affinities for respective metal binding positions. Similarly, the chain B-bound two calcium ions have their B-factor of 37.63 and 47.04 Å^2^. However, for the bound substrate, 2′AMP, the B factors for A and B protomers were found to be similar, 38.87 and 37.04 Å^2,^ respectively. Interestingly, in both chain-A and chain-B, the active site mobile loops have adopted an open conformation similar to chain-B of the IPD-bound structure, and both chains of the apo PaIMPase structure. The absence of the third metal ion in PaIMPase-Ca^2+^-2′AMP complex may account for the open conformation of the active site mobile loop.

**Figure 3:**
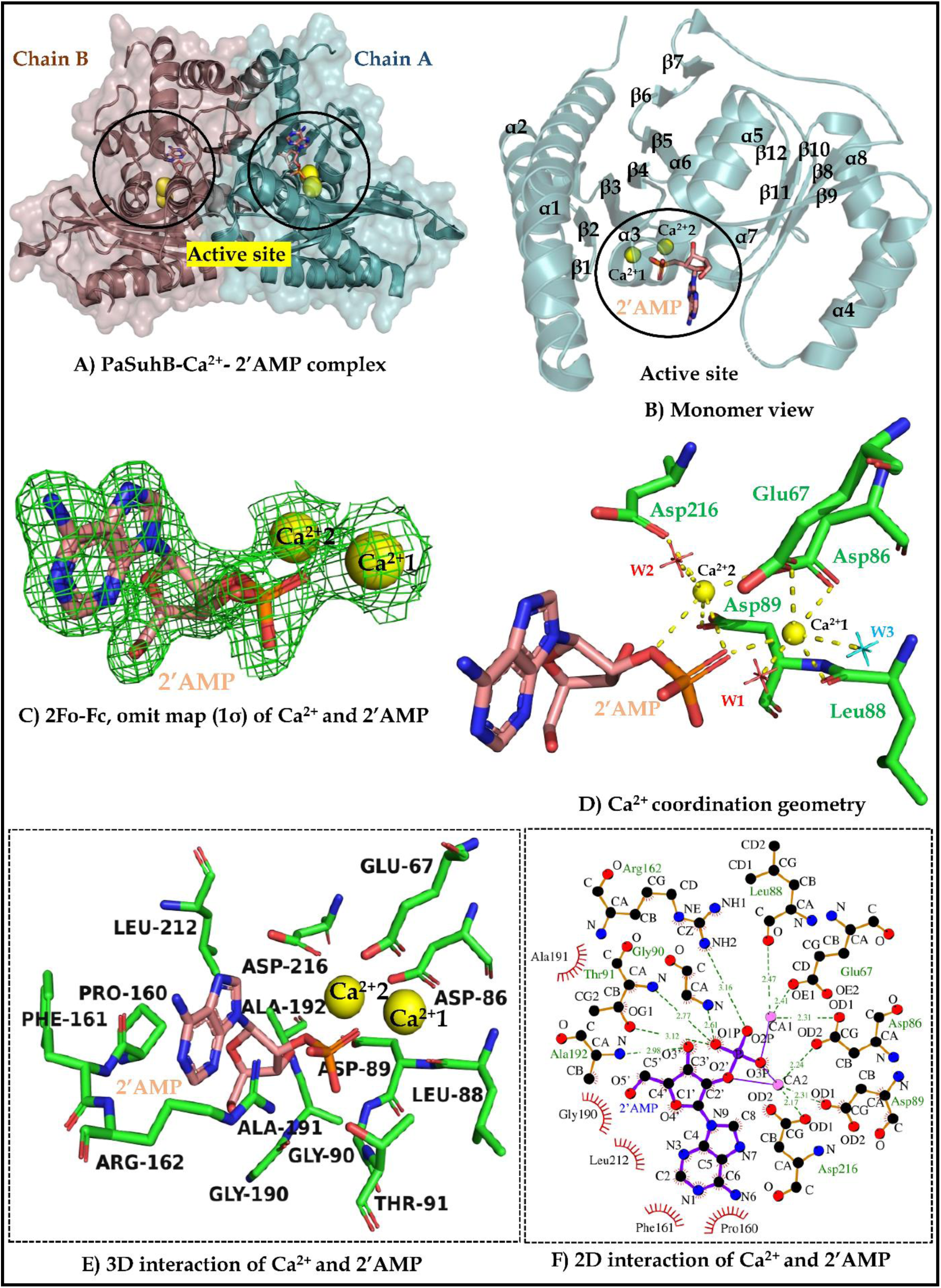
Crystal structure and substrate, metal interaction profiles of PaIMPase-Ca^2+^-2′AMP complex: A) Represents the dimeric PaIMPase/Ca^2+^/2’AMP structure, two circles indicate the active site at each monomer of the dimeric PaIMPase structure. B) Represents the secondary structural architecture of one monomer (Chain-B) showing active site-bound ligands [chain-A has a close resemblance to chain-B (RMSD: 0.093 Å) and is not shown here for pictorial clarity], active site guarding mobile loop and active site proximal a4 helix. C) Represents the 2Fo-Fc omit map (contoured at 1σ) of the PaIMPase active site-bound ligands (Ca^2+^ and 2’AMP). D) Represent the coordination geometry of the bound Ca^2+^ ions at the active site of chain-B. Water molecules involved in the formation of the coordination sphere of the bound metals are indicated by cyan colored asterisks, whereas a couple of water molecules, which have indispensable roles in metal coordination as well as substrate catalysis, are shown by red asterisks. E) and F) Show 3D and 2D interaction profiles of the active site-bound ligands. Throughout the figure, spheres colored in yellow represent bound Ca^2+^ ions, and the bound substrate 2′AMP has been represented by chocolate-colored sticks.

### 3.4. Crystal structure of PaIMPase complexed with substrate transition state analogue

Our attempts to get a cocrystal of PaIMPase complexed with a plausible substrate transition state analogue, tungstate, failed several times. Therefore, the single diffraction quality crystals of PaIMPase in its apo form were used to soak a high concentration of sodium tungstate in the presence of Mg^2+^ (100 mM MgCl_2_.6H_2_O and 20 mM of sodium tungstate), and glycerol at a concentration of 10 % (V/V) was used as a cryoprotectant. These attempts also failed many times due to crystal melting and precipitation of mother liquor in the presence of metal and tungstate together. Finally, PaIMPase apo crystals formed in 0.1 M sodium acetate, pH 5.0, and 20% (V/V) PEG 3350 (**Supplementary Figure S1A2**) were carefully picked and soaked in the mother liquor supplemented with 100 mM MgCl_2_. 6H_2_O, 20 mM Sodium tungstate trihydrate, and 10% (V/V) glycerol for a very short period of time (3 minutes), which yielded good results. The crystals diffracted up to 2.55 Å resolution. A typical diffraction frame of this crystal is shown in **Supplementary Figure S1B2**. After the full set of data collection, the diffraction data of this crystal was indexed and scaled in the P2_1_2_1_2_1_ space group with unit cell dimensions: a=64.59, b=89.483, c=98.137, α=β=γ=90^ο^. The asymmetric unit of the crystal contains a single dimer of PaIMPase with a Matthews coefficient and corresponding estimated solvent content of 2.36 and 47.99 %, respectively (**Figure 4A**). The diffraction data could be rationally processed up to 2.55 Å following all the quality assurance parameters of data reduction (Table 1). The atomic coordinates of the structural model were refined until the R and R_free_ values were merged up to 0.2263 and 0.2849, respectively. The data reduction statistics and the stereochemical, geometric parameters of the final refined model are in good agreement with the crystallographically accepted values (Table 1 and **Supplementary Figure S2A2**). The overall dimeric structure of PaIMPase/Mg2+/GTG complex is stabilised by the formation of eleven hydrogen bonds and six salt bridges, thus accounting for the dimer interface area of ∼1730.9 Å^2^ (**Supplementary Figure S2B2**).

**Figure 4:**
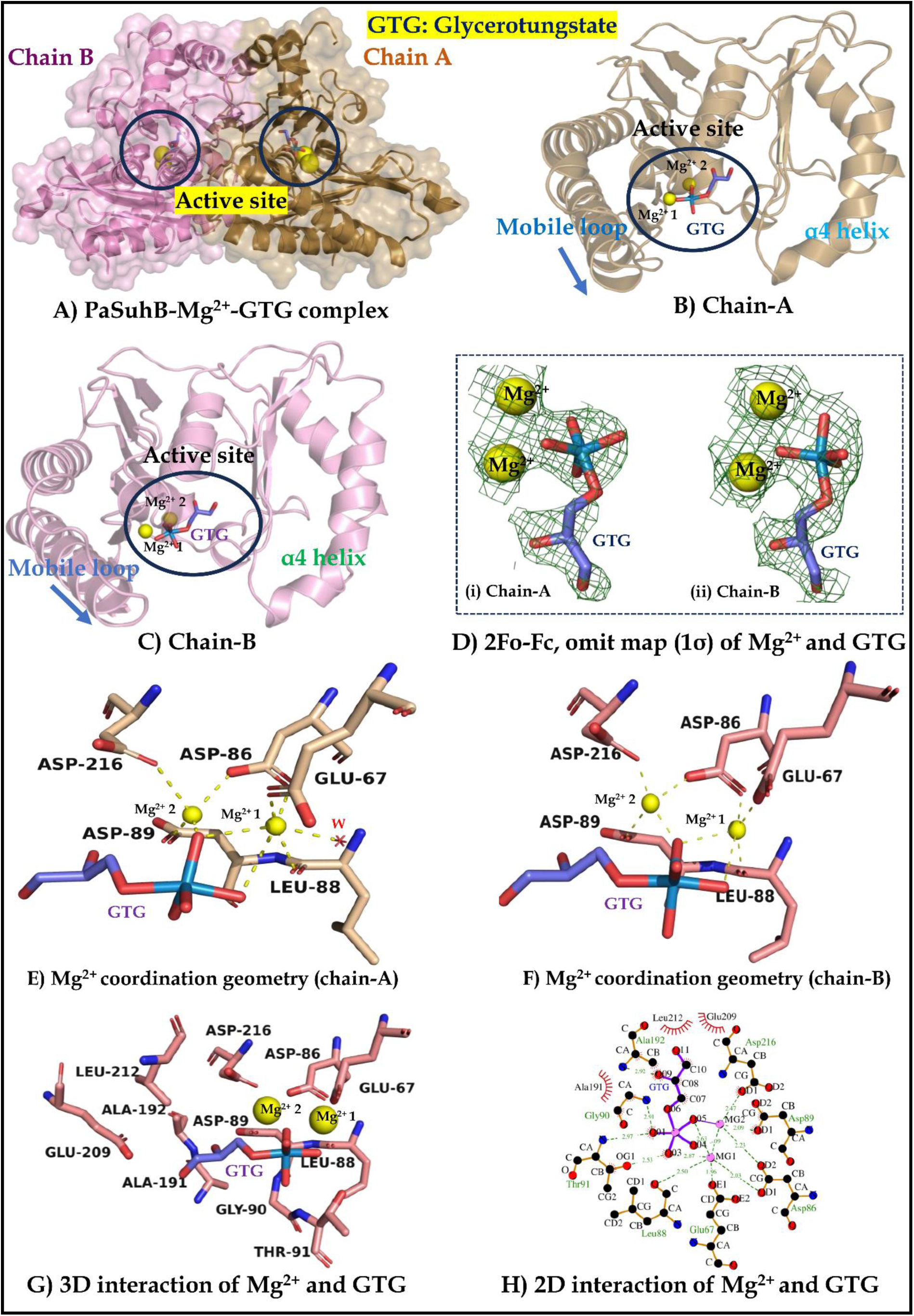
Crystal structure and substrate transition state analogue and metal interaction profile of PaIMPase-Mg^2+^-GTG complex. A) Represents the dimeric view of the PaIMPase-Mg^2+^-GTG complex structure, with two circles indicating the active site at each monomer. B) and C) represent the secondary structural architecture of each monomer, showing active site-bound ligands, mobile loop and a4 helix. Figure D) represents the 2Fo-Fc omit map (contoured at 1σ) of the active site-bound ligands. Figures E) and F) represent the coordination geometry of the Ca^2+^ ions at the active site of chain-A and chain-B, respectively. Water molecules involved in the formation of the metal coordination sphere are indicated by red colour. Figures G) and H) show 3D and 2D interaction profiles of the bound metal ions and the GTG, respectively. Throughout the figure, spheres colored in yellow represent bound Mg^2+^ ions and the bound substrate transition state analogue, GTG, has been represented by blue-colored sticks.

The dimeric and monomeric architectures of the transition state analogue-bound PaIMPase crystal structure are very similar to the substrate-bound structures (Cα RMSD: 0.293 and 0.301 Å with IPD and 2′AMP-bound PaIMPase structure at the dimeric level) with the classical N-αβαβα-C penta-layered overlay of highly conserved secondary structural organisation (**Supplementary Figure S4**). Both the monomers of PaIMPase-Mg^2+^-GTG dimeric structure are very similar **(Figure 4B-C)** with each other, with an overall Cα RMSD value of 0.186 Å (**Supplementary Figure S5**). The average B-factor analysis indicates that the transition state analogue-bound crystal structure of PaIMPase (average B-factor 47.01) is slightly less stable than its two substrate-bound forms (average B-factor 27.64 for IPD-bound PaIMPase and 41.41 for 2′AMP-bound PaIMPase). The difference in the B factors amongst these structural forms of PaIMPase may either stem from the highest possible resolution ranges up to which the diffraction data of the corresponding crystal frames could be processed. However, for the transition state analogue-bound crystal structure of PaIMPase, the drifting of the B factors to a higher side is quite expected, considering the formation of penta-coordinated trigonal bi-pyramidal tungstate, which mimics the transient, high-energy state of the pentavalent phosphorus of the bound PaIMPase substrate. During the initial cycles of structure refinement, the presence of huge difference density peaks (12.00 and 10.75 σ at the active site of chain-A and chain-B, respectively) led us to model two bound metal ions (Mg^2+^) and one tungstate (WO_4_^2-^) ion at the active site catalytic cleft of PaIMPase. Interestingly, after initial fitting and refinement of two metal (Mg^2+^) ions and the tetrahedral WO_4_^2-^ ion inside the difference density peak at the active site of PaIMPase, it was quite evident that the shape of the initially fitted tetrahedral WO_4_^2-^ was not properly fitting inside the obtained difference density peak.

Moreover, an additional conjoined difference density blob remained empty even after the initial fitting of two metal (Mg^2+^) ions and the tetrahedral WO_4_^2-^ ion inside the obtained difference density peak. The additional blob appeared at the distal site of the metal binding cleft and was found to be partly occupying the inositol ring binding site of the PaIMPase active site cavity (as it was observed in PaIMPase/Ca^2+^/IPD complex). Intriguingly, glycerol, a small polyol which was present at high concentration [10% (V/V), which is equivalent to 1.37 M] as a cryoprotectant during the crystal soaking experiment, could be efficiently modelled and successfully refined inside the additional electron density blob neighbouring the central WO_4_^2-^ ion. Glycerol, a small polyol, at high concentration may partially mimic one of the IPD hydrolysis reaction products, inositol, which is also a cyclic polyol. Based on these findings, we also tested the product inhibitory capacity of glycerol against the inositol monophosphatase activity of PaIMPase. We found that at high concentrations [10% (V/V) and beyond], glycerol can also act as a weak inhibitor of the enzymatic hydrolysis of IPD (**Supplementary Figure S6**). Once again, the modelled glycerol moiety was too close (<1.8 Å) to the earlier modelled central WO_4_^2-^ ion. In this situation, we postulated that the proximity of the O1 atom of the modelled glycerol with the proton-withdrawing carboxylate side chain of Asp89 (3.3 Å) and the potential Lewis acid Mg^2+^2 ion (3.2 Å) may result in the formation of activated glycerate anion, which may further attack the nucleophilic centre of the nearby tungsten atom of the WO_4_^2-^ ion, through a SN2 mechanism; thus forming a trigonal bi-pyramidal pentavalent glycero-tungstate adduct (GTG) where the central Tungsten (W) would adopt a pentavalent coordination state. Importantly, the trigonal bi-pyramidal conformation of the pentavalent tungstate adduct (GTG) closely mimics the trigonal bipyramidal transition state of the phospho-monoester hydrolysis, which results from the enzymatic catalysis of many important cellular phosphatases, including inositol monophosphatase (Peck *et al*., 2016). However, unlike the extremely short-lived trigonal bipyramidal transition state of the phospho-monoester hydrolysis, the analogous pentavalent tungstate adduct is relatively stable in solution (Wolfenden, 1969). Subsequent structural modelling, fitting and refinement of the glycero-tungstate adduct (GTG) along with two neighbouring Mg^2+^ ions, finally satisfied the difference density blobs that appeared at the active site of PaIMPase crystal structure soaked with 100 mM MgCl_2_.6H_2_O, 20 mM Sodium tungstate. 3H_2_O and 10% (V/V) glycerol.

Hence, the active sites of PaIMPase dimer are occupied by two molecules of glycerotungstate (one at each active site) and four molecules of Mg^2+^ (two at each active site) (**Figure 4A-C**). The trigonal bipyramidal geometry of the bound GTG at the active site, which mimics the substrate transition state of the IMPase catalysis, was in good agreement with the shape of the electron density blob and the structural parameters like b-factor, density fit analysis, and ligand geometry. As mentioned earlier, our initial trials to fit and refine the tetrahedral WO_4_^2-^ ion in the same difference density blob resulted in a higher b-factor (>100 Å^2^) of the fitted WO_4_^2-^ ion with simultaneous appearance of positive (+*mF_o_*-*DF_c_*) and negative (-*mF_o_*-*DF_c_*) difference density blobs surrounding the ligand. Even optimising the occupancy of the initially fitted WO_4_^2-^ ion did not solve this problem, indicating the fitted WO_4_^2-^ ion is not the right ligand. Although the finally modelled TBP conformation of Glycerotungstate adduct differs from the classical TBP, where equatorial-equatorial bonds should have an angle of 120°, axial-equatorial bonds should have an angle of 90°, and the axial-axial bond should have an angle of 180°. Due to the stereochemical torsions at the PaIMPase active site to favour successful ligand -protein interactions, the tungstate of GTG adduct attains a partially distorted TBP geometry. However, the partially distorted TBP geometry of tungstate is also reported previously in an alkaline phosphatase-bound crystal structure (PDB ID: 5C66). The 2Fo-Fc omit map of Mg^2+^ and GTG, contoured at 1σ, bound at each active site of PaIMPase, is shown in **Figure 4D**. Each of these bound metal ions and the substrate transition state analogue, GTG molecules, could be refined to full (100%) occupancy.

At the active site of chain A monomer, one of the Mg^2+^ ions (Mg^2+^ 1) attains the tetrahedral coordination geometry by interacting with the three highly conserved metal binding residues (Asp86, Leu88 and Glu67) of PaIMPase, one bond from GTG (O5), while the other Mg^2+^ ion (Mg^2+^ 2) attains the tetrahedral geometry by interacting with three conserved metal binding amino residues (Asp86, Asp89 and Asp216) and one bond from GTG molecule(O5) (**Figure 4E**). Contrary to this, at the active site of chain B, metal ion (Mg^2+^ 1) attains a distorted octahedral geometry, while Mg^2+^ 2 attains a tetrahedral geometry with a similar type of interaction as at chain A (**Figure 4F and Supplementary Table 1).** Overall residues from the active site cavity, which are involved in the Mg^2+^ and GTG interaction, are as follows: Glu67, Asp86, Leu88, Asp89, Gly90, Thr91, Ala191, Ala192, Glu209, Leu212 and Asp216 (**Figure 4G-H**). However, the catalytically important water molecules, W1 and W2, which play crucial roles in substrate hydrolysis and product release, were not detected at the active site of the PaIMPase-Mg^2+^-GTG-bound structure. Other than the highly conserved metal-binding residues, the peptide nitrogen of Gly90, Ala192, and the side chain β-hydroxyethyl group of Thr91 stabilise the transition state analogue (GTG) by forming hydrogen bonds with its O1 and O9 atoms. The two magnesium ions bound to the A chain of PaIMPase/Mg^2+^/GTG complex have B-factors of 45.66 and 44.30 Å^2,^ while the two magnesium ions bound to the B chain have B-factors of 47.44 and 43.92 Å^2^. Moreover, the transition state analogue GTG, bound at the chain A and chain B of PaIMPase/Mg^2+^/GTG complex, have a B-factor of ∼40.00 and ∼42.00 Å^2,^ respectively. These B-factor analyses of bound metals and GTG at the PaIMPase active site may suggest that both the monomers of PaIMPase-Mg2+-GTG structure are very similar in their structural disposition, stability in terms of interaction profiling and ligand binding potentials. Most importantly, in none of the PaIMPase active sites of the PaIMPase/Mg^2+^/GTG complex, the traces of the third activating metal ion (Mg^2+^) could be found. Concomitant to this finding, in this structure, the active site mobile loop was also found to be disengaged from the catalytic centre of PaIMPase. Therefore, although the crystal structure of PaIMPase/Mg^2+^/GTG complex, for the first time, represents the plausible structure of the phospho-monoester substrate transition state (through its structural analogue), however, due to the absence of the third activating metal ion and the lack of proximity of the active site mobile loop, the true structural snapshot of the PaIMPase active site during the presence of substrate transition state remained elusive. However, this limitation can be overruled by structurally superimposing PaIMPase/Mg^2+^/GTG complex with the A chain of PaIMPase/Ca^2+^/IPD complex, which demonstrates the immediate precatalytic state of the bound IPD substrate in the presence of three metal ions (Ca^2+)^. Therefore, the comprehensive catalytic mechanism of PaIMPase can be obtained with structural superimposition of different ligand-bound states of the protein, which represent static snapshots of its different catalytic events during substrate hydrolysis.

### 3.5 Crystal structure of PaIMPase complexed with Products myo-inositol (MI) and Phosphate (PO_4_^3-^)

Obtaining the crystal structure of PaIMPase in complex with products (MI and PO_4_^3-^) in the presence of the activator metal Mg^2+^ proved highly challenging, with many failed attempts to co-crystallise the protein in the presence of Mg^2+^ (or Ca^2+^), MI, and PO_4_^3-^. Despite many trials optimising protein-to-product (s) concentration, the co-crystallisation trials were unsuccessful. Even soaking of apo-PaIMPase crystals with high concentrations of Mg^2+^, MI and PO_4_^3-^ was not effective to trace product molecules at the PaIMPase active site catalytic cleft. After several such failed attempts, we replaced the metal Mg^2+^ with Ca^2+^ and soaked the apo-PaIMPase crystals (**Supplementary Figure S1A3**) in the crystallisation mother liquor supplemented with 200 mM CaCl_2_·2H_2_O and 100 mM of each MI and PO_4_^3-^. A typical diffraction frame of this crystal is shown in **Supplementary Figure S1B3**. The diffraction data of this crystal were indexed and scaled in the P2_1_2_1_2_1_ space group with unit cell dimensions, a=65.006, b=88.904, c=98.878, α=β=γ=90^ο^. The asymmetric unit of the crystal contains a single dimer of PaIMPase with a Matthews coefficient and corresponding estimated solvent content of 2.38 and 48.41 %, respectively (**Figure 5A**). The atomic coordinates of the structural model were refined until the R and R_free_ values were merged up to 0.1847 and 0.2413, respectively. The X-ray diffraction data collection statistics and the refined model’s quality assessment parameters fall within the expected values (Table 1 and **Supplementary Figure S2A3**). The active sites of the PaIMPase dimer were found to be occupied by one molecule of MI (one in each dimer), one Ca^2+^ ion (one in each dimer), one PO_4_^3-^ ion (found in chain B of PaIMPase dimer) and one molecule of acetate ion (found in chain A of PaIMPase dimer; binding at the same site of PO_4_^3-^ as in chain B). The overall dimer of PaIMPase/Ca^2+^/MI/PO_4_^3-^ complex is stabilised by 18 hydrogen bonds and 11 salt bridges formed between both interacting monomers with a dimerisation interface area of 1873.3 Å^2^ **(Supplementary Figure S2B3)**. Similar to PaIMPase -apo, -substrate-bound, and -transition state analogue-bound structures, the monomeric structure of each dimer of PaIMPase/Ca^2+^/MI/PO4^2-^ complex (**Figure 5B-C**) preserves its conserved penta-layered architecture of alternating structural layers of alpha helices and β-sheets (αβαβα). Both monomers are very similar to each other with respect to secondary structural arrangements and orientation of the active site mobile loop (RMSD: 0.155 Å), except for the type of ligands bound at the active site (**Supplementary Figure S7**). Each of the active sites of PaIMPase contains a single Ca^2+^ at the classical second metal binding site, one molecule of MI and one molecule of either acetate (at chain-A) or PO_4_^3-^ (at chain-B) (**Figure 5B-C**). The acetate and PO4^3-^ are found to occupy a very similar location in the active site catalytic cleft of the respective PaIMPase monomer. The 2Fo-Fc omit map (contoured at 1σ) of Ca^2+^ MI, and PO_4_^3-^, bound at the active site of the B chain of PaIMPase dimer, is depicted in **Figure 5D**. Each of these bound metal ions and the substrate molecules could be refined to full (100%) occupancy. At both chains, the Ca^2+^ attains the distorted octahedral geometry by interacting with Asp86, Asp89, Asp216, O2 of MI, and two water molecules at chain A and with Asp86, Asp89, Asp216, O2 of PO_4_^3-^ and two water molecules at chain B, respectively (**Figure 5E-F and Supplementary Table 1**). The average B-factor analysis indicates that the PaIMPase/Ca^2+^/MI/PO_4_^3-^ complex structure (with average B-factor 42.32 Å^2^) has a similar average B-factor to the PaIMPase/Ca^2+^/2′AMP structure (41.41 Å^2^) and the apo-PaIMPase structure (40.31 Å^2^), while higher than the PaIMPase/Ca^2+^/IPD structure (27.64 Å^2^). Importantly, the PO_4_^3-^ ion present in the B chain of PaIMPase/Ca^2+^/MI/PO_4_^3^ is shifted from its original catalytic position by 3 Å, as it was evident from PaIMPase/Ca^2+^/IPD and PaIMPase/Ca^2+^/2’AMP complex structures. Nonetheless, the analysis of stereochemistry around the central phosphorus atom of the bound PO_4_^3-^ ion suggests the imminent inversion of stereochemistry of the bound PO_4_^3-^ in PaIMPase/Ca^2+^/MI/PO_4_^3-^ structure compared to that found in PaIMPase/Ca^2+^/IPD or PaIMPase/Ca^2+^/2′AMP complexes. The essential absence of the 1^st^ and 3^rd^ metal ions, the disengagement of the active site mobile loop, as well as the positional shift of the bound free PO_4_^3-^ ion from its catalytic position, may suggest the B chain of PaIMPase/Ca^2+^/MI/PO_4_^3-^complex represents a collapsed product complex. Moreover, the presence of the acetate ion in the same place as PO_4_^3-^ at the Ca^2+^ supplemented A chain of PaIMPase active site explains why the initial co-crystallisation and soaking experiments with lesser concentrations of PO_4_^3-^ did not work in the presence of the 100 mM acetate in the crystallisation mother liquor. Overall, 3D and 2D interaction profiles of the PO_4_^3-^ and MI indicate that the active site amino acid residues, which are involved in interactions, include Asp86, Asp89, and Asp216 and Asp89, Gly90, Thr91, Arg188, Gly190, Ala191, Ala192, and Glu209, respectively (**Figure 5G-H**). The calcium (Ca^2+^ in purple colour 2D figure 5.6H2) interacts with three conserved amino acid residues Asp86, Asp89 and Asp216. This Ca^2+^-coordinating water molecule supports the release of products MI and PO_4_^3-^ from the active site. The chain A and chain B-bound Ca^2+^ have B-factors of 50.09 and 46.22 Å^2,^ respectively. Similarly, the chains A and B bound to MI have their B-factor of 50.99 and 46.96 Å^2^, respectively. The acetate at chain-A has a B-factor of 74.36 Å^2,^ and the PO_4_^3-^ at chain-B has 66.86 Å^2^. Interestingly, in both chain-A and chain-B of PaIMPase protomers, both the active site mobile loops have adopted an open conformation similar to chain-B of IPD-bound structure and both chains of the apo PaIMPase structure, thus not covering the active site, which favours the release of products from the active site and availability of the empty active site for a subsequent round of metal and substrate binding for catalysis. The interaction of MI at the active site involves Asp89, Gly90, Arg188, Gly190, Ala191, Ala192 and Glu209, out of which Gly90, Arg188 and Ala192 form hydrogen bonds while the rest of the residues form hydrophobic interactions. The second released product, PO_4_^3-^, was found to interact with the Ca^2+^ only, which suggests a weak interaction (also justifies the high B factor of bound PO_4_^3-^) and indicates that the products are about to leave the PaIMPase active site.

**Figure 5:**
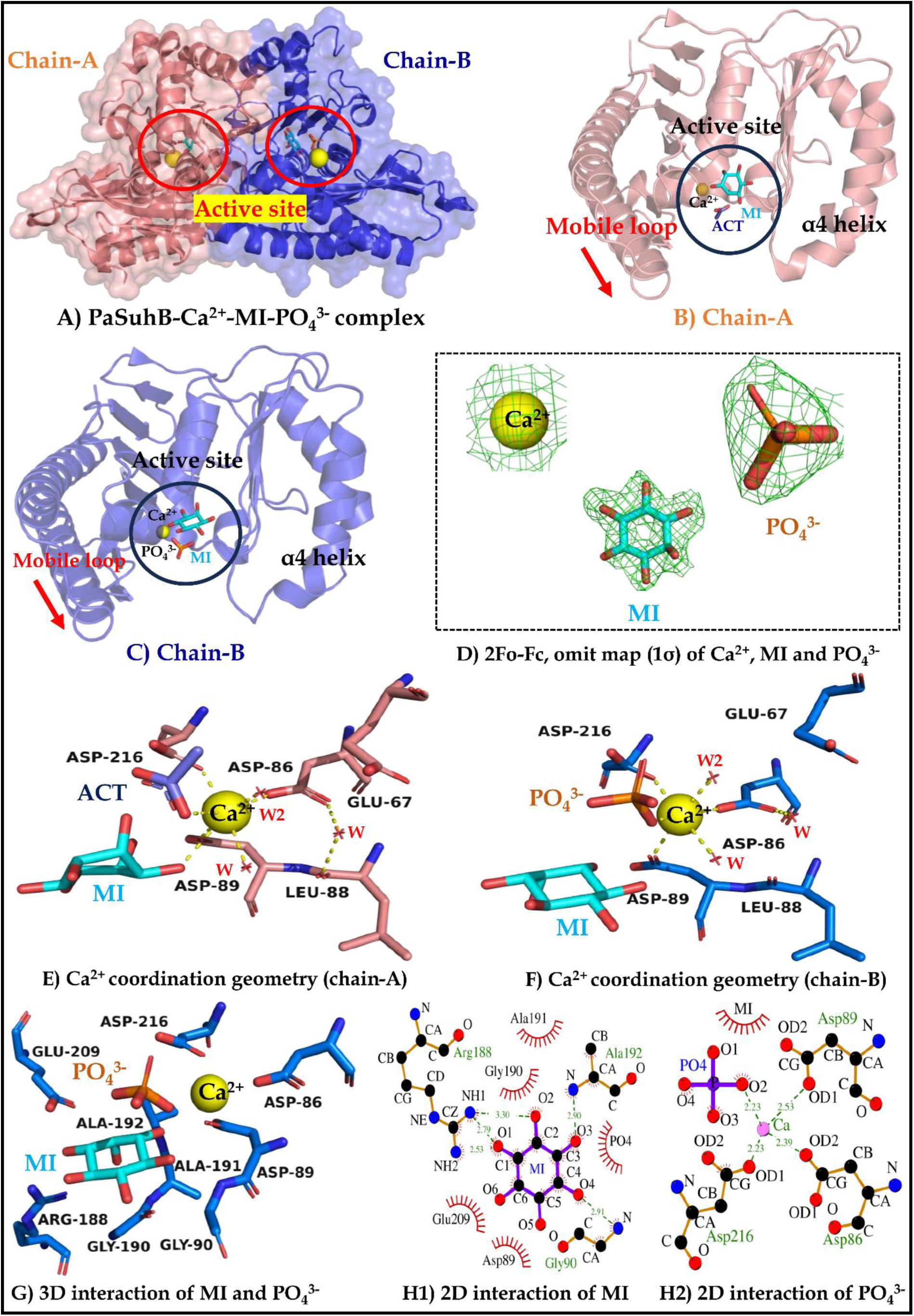
Crystal structure and metal, reaction product(s) interaction profile of PaIMPase-Ca^2+^-MI-PO_4_^3-^complex: A) Represents the dimeric view of the structure, two circles indicate the active site at each monomer. B) and C) represent the secondary structure view of each monomer showing active site bound ligands, mobile loop and a4 helix. Figure D) represents the 2Fo-Fc omit map of the active site-bound ligands contoured at 1σ. Figures E) and F) represent the coordination geometry of the Ca^2+^ ions at the active site of chain-A and chain-B, respectively. Waters responsible for metal coordination are indicated by red colour fonts (W). Figures G) and H) show 3D and 2D interaction profiles of active site-bound ligands, respectively. Throughout the figure, yellow spheres represent Ca^2+^ and cyan, orange and royal blue sticks represent MI, PO_4_^3-^ and ACT, respectively.

### 3.6 Mechanism of metal-assisted substrate binding, active site mobile loop closure and activation of the nucleophilic water

This manuscript aims to decipher the molecular mechanism of enzyme catalysis of IMPases, taking PaIMPase protein as the representative of the IMPase family of enzymes. In this context, we have applied macromolecular crystallography to obtain the structural coordinates of the PaIMPase protein in complex with its substrate (2′AMP), transition state analogue (GTG) and products (MI and PO_4_^3-^). The whole purpose of solving these crystal structures was to capture the snapshots of the different molecular events happening sequentially at the PaIMPase active sites, from the empty enzyme pocket to product release during the course of catalytic events. Since we already have the solved crystal structures of the major events, including apo PaIMPase representing empty active sites and bound with substrates, transition state, and products. We have two crystal structures of substrate-bound forms of PaIMPase dimers, one with IPD and the other with 2′AMP. Four substrate-bound monomers of the protein represent the different stages of the metal and substrate binding. The two monomers of PaIMPase are bound with two metals (Ca^2+^) and 2′AMP (in PaIMPase/Ca^2+^/2’AMP complex); however, in PaIMPase/Ca^2+^/IPD complex, one monomer is occupied with two metals and IPD and the other one is occupied with three metals and IPD. We also have the structural information of two PaIMPase monomers bound with two different substrates (in the presence of two Ca^2+^ ions), which is going to provide significant insights about the plausible substrate-based specific alteration in the conformation of the catalytic site (**Supplementary Figure S8**). It is expected that through these structural snapshots, along with the previously determined structural snapshots of other IMPase orthologues, we will be able to assess the most plausible comprehensive picture of sequential ligand (metal, substrate, substrate transition state, and products) binding and debinding at the active site catalytic cleft of the enzyme, pertinent to the process of enzymatic catalysis. During the course of analysis, we have given major emphasis on the specific regions of the enzyme (**Figure 6A**), which are directly related to the enzyme catalysis, such as the active site pocket (green coloured pocket with surface representation), active site mobile loop (cyan coloured helices with cartoon representation), and the previously identified substrate specificity regulator, α4 helix and its preceding loop (yellow coloured helix and its preceding loop in cartoon as well as surface representation). To investigate the precise molecular mechanism of metal-induced substrate binding and precatalytic events happening around the catalytic sites, we have performed a comparative study of the four unique monomers **(Supplementary Figure S8**), including PaIMPase apo form (green), PaIMPase bound with two metals and 2′AMP (pink), PaIMPase bound with two metals and IPD (blue), and PaIMPase bound with three metals and IPD (cyan). The very first thing we have investigated is the impact of the number of metals on the 3D structural disposition of the active site mobile loop (**Figure 6B**) and the individual amino acid residues from the loop (**Figure 6C**), which makes a significant contribution to the substrate binding and facilitates the precatalytic processes.

**Figure 6:**
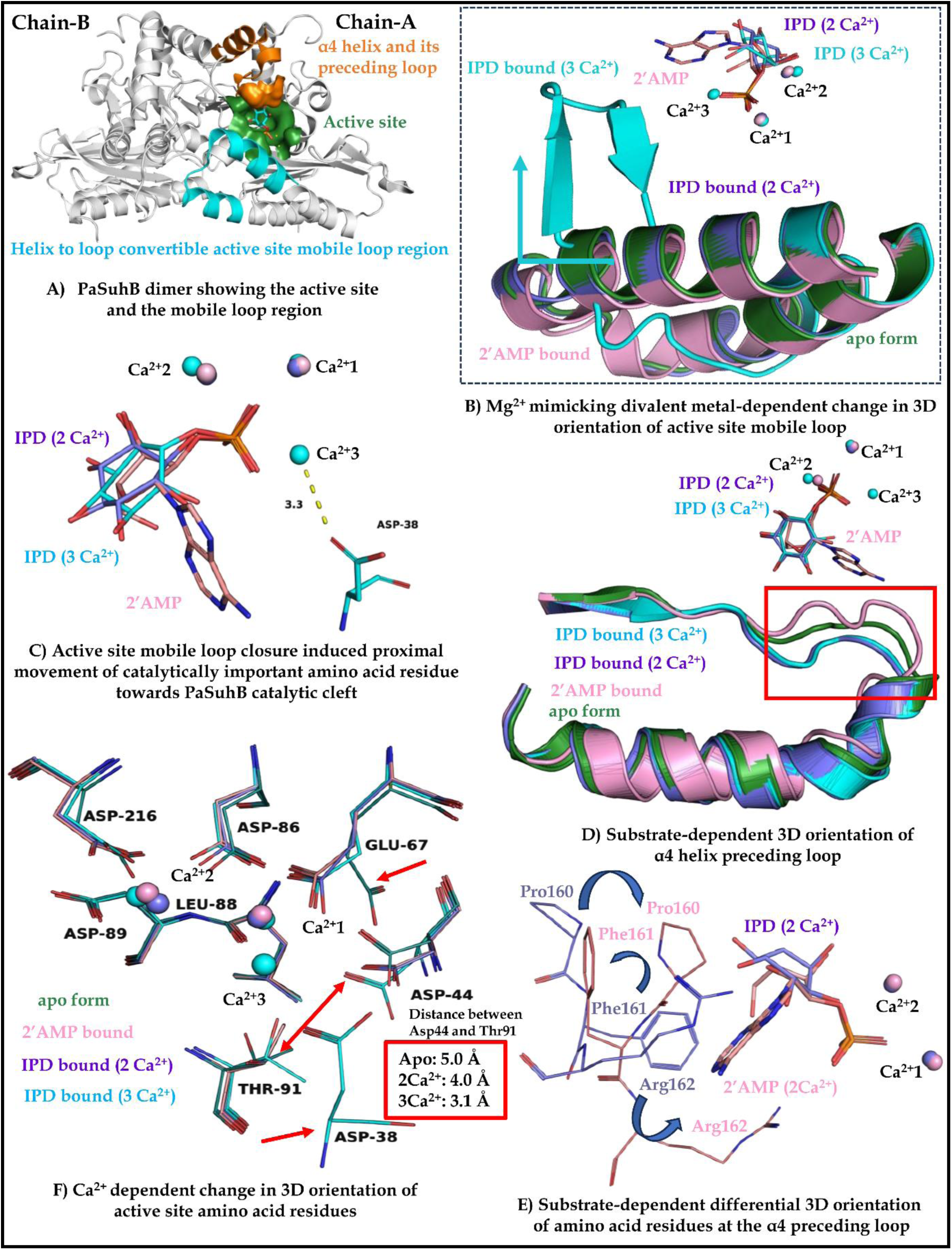
Metal and substrate-induced conformational changes in the active site of PaIMPase. A) PaIMPase-Ca^2+^-IPD bound structure showing active site (green), mobile loop (cyan) and α4 helix and its preceding loop (orange) B) Effect of 3^rd^ Ca^2+^ binding on the 3D structural disposition of the active site mobile loop, C) Catalytically important amino acid residues that orient in the close vicinity of the PaIMPase active site due to 3^rd^ Ca^2+^ binding and concomitant active site mobile loop closure, D) Substrate-induced alteration in the structural disposition of the loop, preceding the α4 helix, majorly altered region is indicated by the red rectangle E) Substrate induced orientation of the majorly altered amino acid residues at the preceding loop of α4 helix, F) Metal induced orientation of the PaIMPase active site amino acid residues. Majorly influenced regions are indicated by red arrows, such as a double-edged arrow that indicates the distance of the Thr/Asp dyad (Thr91/Asp44), which is given in the red box for the respective ligand (metal-substrate) bound structures of PaIMPase. The different ligand-bound states of PaIMPase have been differently colour-coded throughout the presentation, such as Apo structure (Green), IPD bound with two metals (Blue), 2′AMP bound with two metals (Pink) and IPD bound with three metals (cyan).

Interestingly, the results suggest that in the apo-form of PaIMPase, where the active site of the enzyme is empty, the active site mobile loop region of the protein was found to coil back to increase the length of alpha1 and alpha 2 helices by ∼6.5 and 14.5 Å, respectively (Figure 6B green α helix). This structure represents the open conformation of the active site mobile loop of PaIMPase. The coiling back of the active site mobile loop has also been evidenced in the crystal structure of SaIMPase-I (Bhattacharyya *et al*., 2016); however, such conformational change has not been noticed in archaeal orthologues of IMPase (Stieglitz *et al*., 2002). The open conformation of loops was also evidenced in PaIMPase bound with two metals and 2′AMP (Figure 6B pink α helix), PaIMPase bound with two metals and IPD (Figure 6B blue α helix), but the same mobile loop region attains close conformation when the PaIMPase structure is bound with three metals and IPD (Figure 6B cyan α helix), with concomitant helix to loop conformation of this region. These observations strongly suggest that the third metal binding is the hallmark of concomitant closure of the active site mobile loop of PaIMPase. Another important inference that may be extracted from these structures is that the binding of the third metal ion is not necessary for substrate binding, as both of the PaIMPase substrates, 2′AMP (pink) and IPD (blue), are found to bind at the active sites in the absence of the third metal ion. Moreover, it could be speculated from these molecular snapshots that the binding of the two metal ions and the substrate molecule is followed by the third metal ion binding.

We have also evaluated the precise molecular outcomes preceding the active site mobile loop closure event; in this context, we observed that one amino acid residue (Asp38), which was far away in the open loop conformation of PaIMPase, comes into the vicinity of the active site, closer to the third metal ion, when the loop is closed (**Figure 6C**). It is suggested that Asp38 is involved in the correct positioning of the third metal ion at the active site catalytic cleft of PaIMPase. Importantly, the correct positioning of the third metal ion at the active site catalytic cleft of IMPases (including PaIMPase) is mandatory for activation of the water molecule necessary for an inline attack at the nucleophilic phosphorus centre of the active site-bound phosphate monoester, through an SN2 mechanism. Furthermore, we investigated the impact of the number of active sites bound metals on the 3D structural disposition of the substrate specificity-determining α4 helix and its preceding loop. As suggested by the comparative interaction analysis of different metal and substrate-bound forms of PaIMPase, that these regions may support the incoming substrate binding from the distal position of the active site compared to the conventional metal-binding amino acid residues, proximal to the bound metals and the phosphate monoester moiety of the bound substrate; however, there are no major structural alterations of these regions was found to observe due to a change in the number of metals bound to the PaIMPase active site (Figure 6D blue coloured structure with two bound metal ions while the cyan coloured structure is with three bound metal ions). Instead, the structural differences in this region could be observed with different bound substrates (indicated by the red box in **Figure 6D**), an observation aligned with the previously reported role of α4 helix and its preceding loop in the substrate specificity of IMPases (Bhattacharyya *et al*., 2012). These observations prompted us to detect the specific amino acid residues contributing to these structural changes pertinent to substrate binding. We found, in the context of IPD binding, Phe161 was found to be engaged in hydrophobic interaction while the Arg162 was found to a hydrogen bond with O5 of the substrate while the Pro160 was at a non-interacting distance with the substrate; intriguingly, in the 2′AMP bound state of PaIMPase, there is a 180° outward flip of the side chain of the Phe161, which creates the space for a bulky substrate like 2′AMP bearing the purine (adenine) base attached to the 1’ position of the ribose sugar (**Figure 6E**). The resulting orientations of the active site amino acid residues facilitate the binding of 2′AMP via hydrophobic interaction with Pro160 and Phe161 and hydrogen bonding with Arg162. These results infer that PaIMPase demonstrates the conformational plasticity of the active site amino acid residues to facilitate bulky substrate (2′AMP) binding. Next, we analysed the effects of the number of active sites bound divalent metal ions on the 3D structural orientations of the highly conserved metal-binding amino acid residues, as well as the amino acid residues which are directly involved in the water activation and catalysis. Once again, we have employed the same set of different ligand-bound crystal structures of PaIMPase for this investigation, and the results are shown in **Figure 6F**. These comparative studies yield two major structural alterations that were observed sequentially with the apo-, two metal- and the three metal-bound PaIMPase structures. The first observation is the inward orientations of the side chain of Glu67, which is a conserved conventional metal-binding residue that accommodates the first and the third metal ions at the active site of PaIMPase. These two metal ions, in other IMPases, are suggested to activate the water nucleophile, which subsequently is responsible for the hydrolysis of the bound phosphomonoester substrate. Likewise, in PaIMPase, these two metal ions are also expected to exert a similar catalytic role. The differential orientation of the Glu67 side chain in the apo- and metal-bound active site of PaIMPase suggests its crucial role in the positioning and stabilisation of the first and the third metals to aid the subsequent catalytic events (**Figure 6F**). Moreover, the other interesting observation is the relative distance between the Thr91/Asp44 dyad in different metal-bound conformations of PaIMPase. From the earlier studies, it is evident that this conserved Thr/Asp dyad is responsible for extracting the proton from the nucleophilic water molecule, which subsequently attacks the positively charged central phosphorus atom of the bound phosphomonoester moiety of the substrate molecule. The structural superimposition of different ligand (metal and substrate) bound conformation of PaIMPase indicate a gradual decrease in the distance of this dyad as the active site is occupied by subsequent binding of metals and substrate, for example the relative distances in between Thr91 and Asp44 was 5.0 Å in the apo structure, 4.0 Å with two metal and substrate bound form and 3.1 Å with three metals, and substrate bound form of PaIMPase. Finally, in the three metal and substrate-bound immediate pre-catalytic conformation of PaIMPase, these two residues are in the range of hydrogen bonding, which is a prerequisite to snatch a proton (H^+^) from the water nucleophile. These observations signify that the number of metal binding at the PaIMPase active site significantly influences the distance of Thr91/Asp44 dyad to bring them in the hydrogen bonding vicinity only after the binding of the third metal ion, which represents the ideal conditions for proton relay and water activation. Altogether, the comparative structural analysis of different ligand-bound conformations of PaIMPase reveals the molecular basis of metal-assisted substrate binding mechanisms and the sequential catalytic events happening in and around the PaIMPase active site, which represents the precatalytic preparation of the protein for its enzymatic activity.

### 3.7 Mechanism of catalysis and product release

In the previous section, we delineated the molecular events associated with the metals and substrates binding at the active site catalytic cleft of PaIMPase by comparing the different PaIMPase apo and substrate-bound crystal structures. To further accomplish our ultimate goal to unravel the molecular mechanism of enzyme catalysis of PaIMPase, we have performed a detailed structural comparison study of the substrate-bound - precatalytic, transition state analogue-bound mid-catalytic and product-bound - post-catalytic states of PaIMPase. In this context, the monomeric structure of PaIMPase with three Ca^2+^ and IPD bound state represents the plausible precatalytic events (cyan stick and spheres). In contrast, the two Mg^2+^ and GTG bound state of PaIMPase represents the plausible mid-catalytic transition state (magenta stick and spheres). Finally, the Ca^2+^-MI-PO_4_^3-^ bound monomer of PaIMPase-[blue sphere (blue and orange sticks represent MI and PO_4_^3-^ respectively)] represents the plausible post-catalytic states of the protein (**Figure 5.8A and Supplementary Figure S9**). **Figure 7B** indicates the high-resolution snapshot of the precatalytic events supporting the three metal-assisted catalysis of the phosphomonoester substrate, suggested by earlier studies (Lu *et al*., 2012). In the precatalytic phase, all three bound metals satisfy their octahedral coordination geometry supported by the interactions with amino acid residues (through side chain carboxylates as well as peptide carbonyl), phosphomonoester moiety of the bound substrate, and the active water molecules. The coordination sphere of the Ca^2+^-1 consists of O atoms of the Asp86, Leu88, and Glu67, one bond from the phosphomonoester moiety of the substrate and two from neighbouring water molecules. Similarly, the coordination sphere of Ca^2+^-2 contains O atoms of Asp86, Asp89, and Asp216, two from the phosphomonoester moiety of the substrate and one from water. The coordination sphere of the third metal, Ca^2+^-3, consists of the O atom of Glu67, one from the phosphomonoester moiety of the substrate and four from the surrounding water molecules. One water molecule, denoted by W1, is shared in the coordination spheres of Ca^2+^-1 and Ca^2+^-3, and another water molecule, W2, is shared between the coordination spheres of Ca^2+^-2 and Ca^2+^-3 (**Supplementary Table 1**). Earlier studies suggested that to be an activator metal of the IMPases, two parameters are considerably crucial: one is the coordination number of the metal, and the other is the charge-to-mass ratio of the bound metal. We have chosen Ca^2+^ to deduce substrate-bound structural attributes of PaIMPase because it shows the coordination number, coordination geometry and preference for the coordination ligands (soft versus hard) as the same as the Mg^2+^, but Ca^2+^ has a lesser charge-to-mass ratio, which subsequently is not able to polarise water molecules to activate the nucleophile to initiate the substrate hydrolysis. Therefore, Ca^2+^ helps to capture the molecular snapshots of the precatalytic events. The coordination spheres of all three-substrate monoester coordinating bound metal ions in PaIMPase precatalytic complex will render the central phosphorous atom of the phosphomonoester moiety of the substrate an ideal centre for nucleophilic attack. Similarly, the O atom of the W1 is also pulled by Ca^2+^-1 and Ca^2+^-3, causing polarisation in the water (OH-H) molecule. This process is further assisted by the Thr91/Asp44 dyad, which comes in the vicinity of the active site concomitant to the third metal binding. The negatively charged side chain carboxylate anion of Asp44 pulls the polar side chain (1-hydroxyethyl moiety) hydrogen of Thr91, resulting in the partial negative charge on the hydroxyl group of Thr91, which acts as a Lewis base and pulls the hydrogen of the W1 molecules, which was already partially polarised by the metal ions.

**Figure 7:**
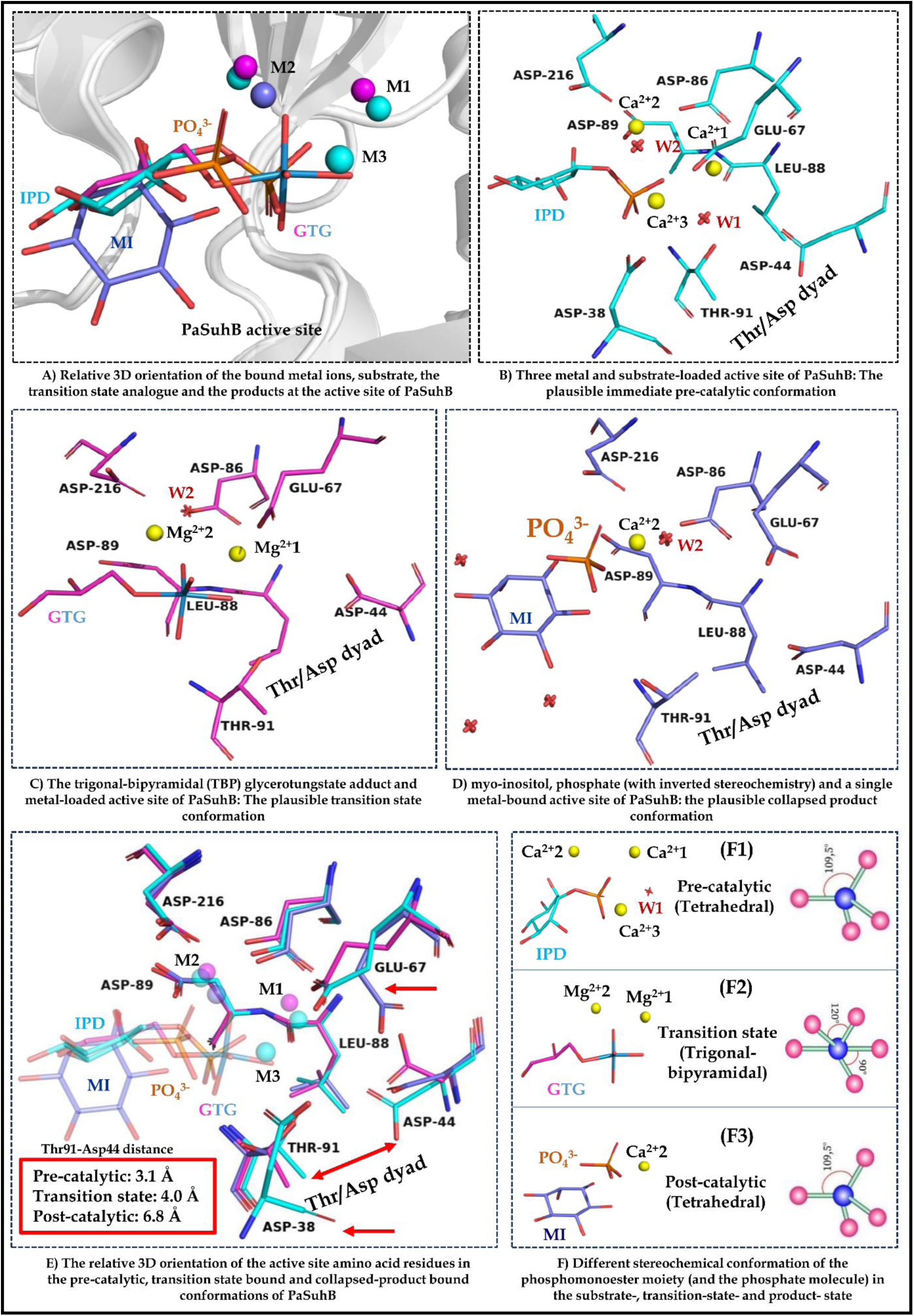
Structural snapshots of different plausible precatalytic, mid-catalytic (transition state) and post-catalytic active site conformations of PaIMPase A) Substrate (IPD), transition state analogue (GTG) and products (MI and PO_4_^3-^) bound at the PaIMPase active site along with the metal ions; M1, M2, and M3 represents metal ions that occupies first, second and third metal positions, B) 3D-Orientation of substrate, metal ions, water molecules and the PaIMPase active site amino acid residues representing the plausible precatalytic event, C) 3D-orientation of transition state analogue, metal ions, water molecules and the PaIMPase active site amino acid residues representing the plausible intermediate state of substrate hydrolysis, D) 3D-orientation of products, metal ions, water molecules and the PaIMPase active site amino acid residues representing a collapsed product state E) Superimposition of precatalytic, mid-catalytic and post catalytic conformations of the PaIMPase active site, residues with major structural fluctuations are indicated by red arrow and the red box shows the relative distances in between Thr91 and Asp44 dyad in different conformational states of PaIMPase, F) Relative stereochemistry of the phosphomonoester moiety (in the substrate and the transition state analogue) and the phosphate molecule (product molecule) around the central phosphorus atom of the substrate, transition state analogue and product at precatalytic, mid-catalytic state and post-catalytic state of PaIMPase, respectively. Throughout the figures, the structure, and their bound metals, water molecules and ligands are represented in identical colours, such as precatalytic (Cyan), transition state analogue (magenta), and post-catalytic (blue). The phosphate and tungstate moieties are shown in orange and royal blue colours, respectively.

Consequently, the formation of reactive hydroxyl groups from the W1 water molecule is imminent, which subsequently attacks the positively charged central phosphorous atom of the phosphomonoester moiety of the bound substrate, thus resulting in the formation of a highly unstable penta-coordinated phosphorane-like trigonal bipyramidal transition state of the central phosphorus atom. Intriguingly, we have solved the crystal structure of the PaIMPase in complex with Mg^2+^ and the pentacoordinate phosphorane-like transition state analogue glycerotungstate (GTG). **Figure 7C** indicates the plausible molecular snapshot of the trigonal bipyramidal intermediate state containing the active site of PaIMPase. This is the first molecular glimpse of IMPase active site loaded with activating metal ions and the pentacoordinate phosphorane mimicking a transition state analogue. Although the PaIMPase/GTG/Mg^2+^ complex structure suggest that the presence of a third metal ion is not necessary once the structure attains the transition state, however, the pentacoordinated structure of tungsten is much more stable than the pentacoordinated phosphorane and actual molecular situation might be quite different while the pentacoordinated phosphorane like transition state is formed, and which might need the presence of the third metal ion. However, the pentacoordinated tungstate adduct truly mimics the phosphorane transition state, as it is evident that the axial oxygen atom lying at the opposite side to the ester oxygen of the glycerotungstate adduct occupies almost the same place as the nucleophilic water molecule (W1) found in the three-metal-based precatalytic complex of PaIMPase. However, the hexa-coordinated first metal ion, which should activate the nucleophilic water (W1) during the precatalytic event, is found to be present in a tetrahedral coordination geometry (Chain-A), suggesting that the octahedral coordination geometry of the activating metal ion may be required only to hold the attacking water molecule, which is not necessary once the transition state is formed. Interestingly, W2, the second water molecule bound to the second and third metal, is still in the close vicinity of the Mg^2+,^ suggesting its role is yet to be accomplished. Previous studies suggested that during post-hydrolysis of the substrate, W2 protonates the leaving first product (the alcohol group; inositolate anion if the inositol monophosphate is the substrate). Thus, to hold the W2 in its proper place, the second metal ion during the transition state still forms a distorted tetrahedral coordination geometry with one additional bond supporting the unhydrolyzed monoester substrate. Moreover, the distance between the Thr91/Asp44 dyad also increase as compare to the precatalytic phase, suggesting that once the water activation is done, all the necessary elements needed for the activation of nucleophilic water are relaxed at the active site at a distance more than that of a stable hydrogen bonding distance. Since the actual phosphorane like transition state is highly energetic and unstable, it needs to be resolved immediately, once it is formed to resolve this, the ester bond connecting the O from MI and the phosphorous centre of the monophosphate breaks resulting in the formation of the two products MI (initially the inositolate ion followed by the formation of inositol upon protonation by W2) and inorganic PO_4_^3-^.

In case of products bound PaIMPase structure, we found that there is only a single metal ion at the second position of the metal binding, in the distorted tetrahedral coordination geometry bound with the side chain carboxylates of three amino acids, Asp86, Asp89, and Asp216, one oxygen of product PO_4_^3-^and one bound water in hydrogen bonding distance (**Figure 7D**). Interestingly, the same structure also indicates that the MI is highly bonded with the neighbouring water molecules, which suggests that the product is in a unionised state and free to leave the active site. Apart from these, at the active site, both products also show very weak interactions. Moreover, the second released product PO_4_^3,^ which also shows an inversion of configuration around the central phosphorus atom compared to its substrate-like configuration found in IPD, shifts from its original position from the active site by ∼3 Å as compared to the IPD-bound and GTG bound structures. The product PO_4_^3-^ binds with only Ca^2+^-2 with a single bond, and MI is already surrounded by water shells. These observations altogether indicate that PaIMPase/Ca^2+^/MI/PO_4_^3-^ complex represents a collapsed product complex, indicating that the active site is about to evacuate all the formed products and will be ready for the next round of catalysis (**Supplementary Figure S12).** Furthermore, the comparative study of the active site residues of precatalytic, transition state and post-catalytic events indicates that the outward movement of Glu67 in product-bound conditions suggests that this orientation of Glu67 represents either an empty active site (in PaIMPase apo structure) or collapsed product state of the post-catalytic event, suggesting that the products are about to leave the active site (**Figure 7D-E**). Likewise, the distance between Thr91/Asp44 dyad and the position of Asp38 is also suggestive of near evacuation of the PaIMPase active site. During the precatalytic phase, the Thr91/Asp44 distance was 3.1 Å. Once the transition state is achieved, this distance is increased to 4.0 Å, which is even further increased after the substrate hydrolysis and product formation, to 6.8 Å (Figure **7E** **red rectangle**). Similarly, the position of the asp38 in 3D space is close to the active site only in the case of the precatalytic phase, while it is far away in the case of the transition state and product-bound crystal structures (**Figure 7D-E**). The coordination geometry of the different catalytic events is given in **Figure 7F**, demonstrating ideal and experimental bond length and orientation of the atoms in the respective geometrical disposition. The ideal tetrahedral geometry represented in **Figure 7F** indicates that all the bonds fall at 109.5° angles from each other. The PO_4_^3-^ group of our substrate also attains the same tetrahedral geometry facing the third metal or activating water w1 (**Figure 7F1**). Once the substrate attains TBP geometry, all three oxygen atoms come in one plane. The ideal angle is given in **Figure 7F2**. When the TBP forms at the enzyme active site, the angle and bond length slightly deviate from the ideal position, which we have explained earlier in this manuscript. This deviation is brought to accommodate the proper binding and successful interactions at the active site. It is believed that when the product is formed post P-O bond breakage, the orientation of the tetrahedral geometry of PO_4_^3-^ seems to be inverted, meaning it is opposite to the third metal or activating water W1 (**Figure 7F3**). In the case of our product-bound structure, we do not observe product geometry inversion significantly. This might be due to the relative position of PO_4_^3-^ was also significantly altered from the PO_4_^3^ of the substrate condition. Moreover, we also investigated the relative orientation of the active site mobile loop during the process of catalysis by comparing the same set of crystal structures. Interestingly, we found that, as in the case of apo and two metal-bound structures, the active site mobile loop is coiled back in an open conformation in the transition state and product-bound states of PaIMPase, except at the precatalytic phase of the third metal-bound PaIMPase (**Supplementary Figure 10**). This observation corroborates that loop closing is only required to activate the nucleophilic water molecule, but not at any other stages of the catalytic event. Collectively, these observations comprehended the detailed mode of catalytic mechanism of PaIMPase and may represent the similar sequential events happening around the active site of the IMPase enzymes.

## 4. Discussion

IMPases are crucial for PI signalling in humans and regulate virulence and pathogenesis in lower-order organisms like bacteria. They are well-known drug targets for bipolar disorder and antibacterial therapy for drug-resistant bacterial pathogens. In this context, tremendous efforts have been carried out to discover novel molecular scaffolds that can effectively counter bipolar disorder and bacterial infections. Since IMPase shares a high degree of 3D fold conservation and conserved active site residues despite very low sequence similarity (**Supplementary Figure S11**), the inhibitors targeting the active site are expected to diminish most of the IMPase enzyme activity. The precise knowledge of the biochemical features and detailed molecular mechanisms of these enzymes can be exploited to design potential inhibitor leads, which can further be developed to eradicate bipolar disorder and bacterial infections. This study aims to decipher the detailed molecular mechanism of PaIMPase by utilising the biochemical and crystallographic features of substrates, substrate transition state analogue and product-bound PaIMPase structures. The IPD and 2′AMP are the only substrates that can be hydrolysed by PaIMPase, suggesting no additional catalytic roles in *P. aeruginosa*, unlike *S. aureus* and thermophilic bacteria, where IMPase also hydrolyses NADP and F-1, 6-BP, respectively (Bhattacharyya *et al*., 2012; Stec *et al*., 2000). Apart from Mg^2+^, Co^2+^, Mn^2+^ and Zn^2+,^ which can also activate this enzyme, whereas Ca^2+^ and Fe^2+^ inhibit its enzyme activity, representing classical IMPase behaviours with respect to metal specificity (Pollack *et al*., 1993). The optimum Mg^2+^ concentration was found to be higher (30 mM) than most other IMPases (< 5 mM), suggesting the PaIMPase can withstand higher salt concentration, likely to hyperthermophilic bacteria. The enzyme kinetic parameters reveal that IPD is the preferred substrate over 2′AMP. The PaIMPase is inhibited by Li^+,^ and higher concentrations of Mg^2+^ demonstrate the classical IMPase characteristic, but again they show the difference in the Mg^2+^ concentration required for inhibition (>100 mM) than the classical IMPase (>5 mM). The K_i_ value of PaIMPase with respect to Li^+^ (2.2 mM) suggests that, like mammalian IMPase, they are one of the Li^+^ sensitive enzymes, unlike Li^+^ insensitive archaeal IMPase (K_i_ 250 mM). To validate these biochemical findings and capture the different snapshots of enzymatic events happening at the active site, the detailed and sequential analysis of the crystal structures of PaIMPase complexed with two substrates, IPD and 2′AMP, substrate transition state analogue (GTG) and products (MI and PO_4_^3-^), along with the apo structure representing an empty active site were critically investigated. Interestingly, the two active sites of the IPD-bound PaIMPase crystal structure demonstrate differential metal binding, one with two and another with three, representing two sequential events during substrate binding. The comparative analyses of apo structure, two metals bound 2′AMP, two metals bound IPD, and three metals bound IPD reveal the mode of metal and substrate binding and some of the interesting findings, such as the role of third metal ions in loop closing. Extensive analysis of the IMPase structures available on PDB indicates that there are many crystal structures with the active site mobile loop in close conformation, irrespective of the number of metals and type of ligands bound at the active sites. Careful examination of these crystal structures suggests that the loop can also be closed by other factors, like the orientation of monomers in dimeric structures, crystal contacts which pushes active site mobile loop into a close conformation, and other interacting partner proteins that facilitate compactness of the dimeric organisation. Apart from all these factors, the sequential metal binding during catalysis facilitates loop closure only after the binding of the third metal. Two metal ions are considered sufficient to stably accommodate the incoming substrates irrespective of IPD or 2′AMP, which is further supported by the analysis of many previously solved crystal structures available on the PDB database (Bone *et al*., 1994). The Asp38 orient in the vicinity of the active site due to loop closure, supporting the previous finding, which states that they are required for the positioning of the third metal and activation of catalytic water (Lu *et al*., 2012; Choe *et al*., 1998). The detailed comparative analyses of IPD and 2′AMP-bound PaIMPase structure indicate that Phe161 was found to regulate the binding of substrates, as there is 180 degrees outward flip when a larger substrate, like 2′AMP, is present. Bhattacharyya et al. have shown the role of α4 helix in substrate specificity of the IMPase family protein, and interestingly, Phe161 is part of the loop preceding α4 helix, which functions as a gatekeeper to adjust the volume of the active site for larger substrates. These observations justify the substrate preference of PaIMPase as IPD over 2′AMP, as additional efforts are required to accommodate 2′AMP, which subsequently lowers the catalytic efficiency.

Moreover, the distance between Asp44/Thr91 dyad lowered to 3.1 Å as the third metal binds, which functions as a H^+^ ion acceptor, facilitating the water (W1) activation with the help of Ca^2+^-1 and Ca^2+^-3 metal ions. Thus, the activated water molecule attacks the positively charged central phosphorus atom, resulting in the formation of the trigonal bipyramidal (TBP) geometry representing the intermediate state of enzyme activity. To our knowledge, this is the first time we have obtained the TBP state intermediate analogue bound crystal structure of the IMPase protein. The absence of the third metal ion in the TBP structure reveals that once the catalysis starts, the third metal ion is not necessary for downstream events. The product-bound crystal structure has many water molecules at the active site surrounding the MI, suggesting that the protonation of the inositolate ion and subsequent product release from the active site. The comparative analysis of precatalytic (three metal-bound IPD), transition state (GTG-bound) and products (MI and PO_4_^3-^-bound IMPase) demonstrates several interesting features like the Glu67, Asp44/Thr91 dyad distance and Asp38 position. The outward orientation of Glu67 during all other phases except at the precatalytic reveal that the third metal ion binding is initiated by the Glu67 residue and governs the entry and exit of the substrate and products, respectively. The Asp38 shows similar characteristics as it is close to the active site only at the precatalytic event, suggesting its role in precatalytic water activation. The distance between Asp44/Thr91 dyad, which was coming close to the active site during the precatalytic event, is going away from the active site during the transition state and post-catalytic phase. The active site mobile loop orientation also suggests that the loop closure is only required during water activation; the rest of the time, it is oriented in an open conformation to facilitate the entry and exit of the substrate and products. The five crystal structures of PaIMPase solved by our group, the previously solved crystal structures of IMPases and the available literature have been extensively collated to decipher the molecular mechanism of IMPase family enzymes.

## 5. Summary and future perspective

Owing to the pathological relevance of IMPases and virulence regulation potentials of PaIMPase, we have cloned, overexpressed, biochemically characterised and solved the crystal structure of PaIMPase enzyme in complex with substrates (2′AMP), transition state analogues (GTG) and products (MI and PO_4_^3-^). These structures, along with the apo and IPD-bound PaIMPase, were critically examined to decipher every plausible molecular event happening at and around the active site of PaIMPase. Crystallographic investigations of the apo structure with substrate-bound forms reveal that the binding of the third metal ion is necessary for the active site mobile loop closure, which subsequently brings the three amino acid residues Asp38 close to the active sites required for metal positioning and water activation. The IPD and 2′AMP binding mechanism demonstrates different orientations of the active site distal amino acid residues Pro160, Phe161, and Arg162. The 180° flip of Phe161 accommodates the binding of 2′AMP at the active site by providing the space for the bulky substrate. The binding of the metal induces the inward orientation of the Glu67, representing the metal ion-occupied active site as compared to the empty active site in the apo PaIMPase structure. The two metal ions binding at the active site catalytic cleft of PaIMPase are sufficient for the binding of substrates, as we also observed in these structures that there is no substrate when there are no metal ions present. This could be apprehended by the fact that the metal binding site of PaIMPase active site is lined with negatively charged amino acid residues, which cannot directly support the binding of the negatively charged phosphate monoester substrates unless the presence of positively charged divalent metal ion adaptors. At the active site catalytic cleft of PaIMPase, the nucleophilic water (W1) was activated to form a catalytic hydroxyl ion due to the positively charged metal ions and negatively charged Thr91, which acts as a Lewis acid in the presence of Asp44. This hydroxyl group attacks the positively charged central phosphorus atom of the bound phosphomonoester substrate to form a phosphorane-like trigonal bipyramidal transition state. This highly unstable transition state leads to the cleavage of the P-O ester bond to form products inositolate and PO_4_^3-^. The first leaving product, inositolate, was protonated by a water molecule (W2), which is present in the coordination sphere of the activating metal ion bound at the second metal binding site. The Thr91/Asp44 dyad is crucial for water nucleophile (W1) activation; the binding of the third metal only brings these two residues close enough to deprotonate and activate the W1, except that condition, these two residues lie too far apart from each other. The phosphomonoester moiety of the bound substrate, transition state and the product PO_4_^3-^ attain tetrahedral, TBP and inverse tetrahedral geometry around the central phosphorus atom during the precatalytic, mid-catalytic and post-catalytic phases of the enzyme activity. Further study indicates that the loop closure is not required once the transition state has been formed, as well as during product formation and release. However, this observation may vary during the actual phosphorane-type transition state formation and resolution into products, as the actual phosphorane-like transition state is much less stable compared to the pentavalent tungstate adduct-based transition state mimic we used herein. The crystal structures presented in this chapter not only shed the molecular mechanism of PaIMPase’s IMPase activity for the first time but also represent the plausible structural attributes of the substrate transition state analogue-bound structure of IMPase class of enzymes for the first time. The most crucial trigonal bipyramidal geometrical state of the substrate transition state bound at the metal-supplemented active site catalytic cleft IMPase is a long-awaited research gap in this field. The comprehensive descriptions of the molecular events happening during the process of catalysis using the single IMPase protein (PaIMPase) are the novel finding discussed in this manuscript. The precise knowledge of the molecular events of catalysis and the coordinates of the enzyme in complex with substrates, transition state analogues, and the products provides significant insights into designing small molecule-based inhibitory leads against this protein to circumvent the pathological consequences of these classes of enzymes.

## Acknowledgements

Special thanks to the Director, IIT Jodhpur, for providing the infrastructure and experimental facility to conduct all the experiments reported in this manuscript. AYURTECH facility at IIT Jodhpur and RRCAT Indore, India, for providing a synchrotron radiation source for protein crystal diffraction and data collection. V.K.Y expresses thanks to the Ministry of Education, Government of India, for providing fellowships to carry out doctoral research. We acknowledge the Copilot AI tool for assisting with the editing and formatting of this manuscript.

## Data availability

The crystal structures and associated information can be accessed from the PDB database with the PDB IDs: 8WDQ, 9WBP, 22HF and 9XCK. All other related data can be found in this manuscript and its supporting information. Apart from that, the corresponding author will provide any data related to this manuscript by following the journal’s guidelines.

