## Supplementary Information for "Unravelling the plausible metal-dependent catalytic mechanism of Inositol monophosphatase ortholog from *Pseudomonas aeruginosa* through the lenses of macromolecular crystallography and enzyme kinetics"

**Table of contents**

| S. No. | Supplementary contents | Page Number |
|---|---|---|
| 1 | S1: Crystallisation and Diffraction Patterns | 2 |
| 2 | S2: Ramachandran plot and dimer interface area | 2 |
| 3 | T1: Interaction profiles of divalent cations at the active sites | 3 |
| 4 | S3: Monomers of 2′AMP bound crystal structure | 4 |
| 5 | S4: All crystal structures showing conservation at dimeric levels | 4 |
| 6 | S5: Monomers of GTG bound crystal structure | 5 |
| 7 | S6: Glycerol-based PaIMPase activity inhibition | 5 |
| 8 | S7: Monomers of MI and $PO_4^{3-}$-bound crystal structure | 6 |
| 9 | S8: Overlapped monomers of apo and substrate-bound PaIMPase | 6 |
| 10 | S9: Overlapped monomers of pre-catalytic, transition state and post-catalytic state representing PaIMPase | 7 |
| 11 | S10: Representation of active site mobile loop orientations during the process of catalysis | 7 |
| 12 | S11: Structure and sequence comparison of human and *P. aeruginosa* IMPase | 8 |
| 13 | S12: Comparative 3D positioning of the $PO_4^{3-}$ from substrate, immediate product, and collapsed product-bound crystal structures | 9 |

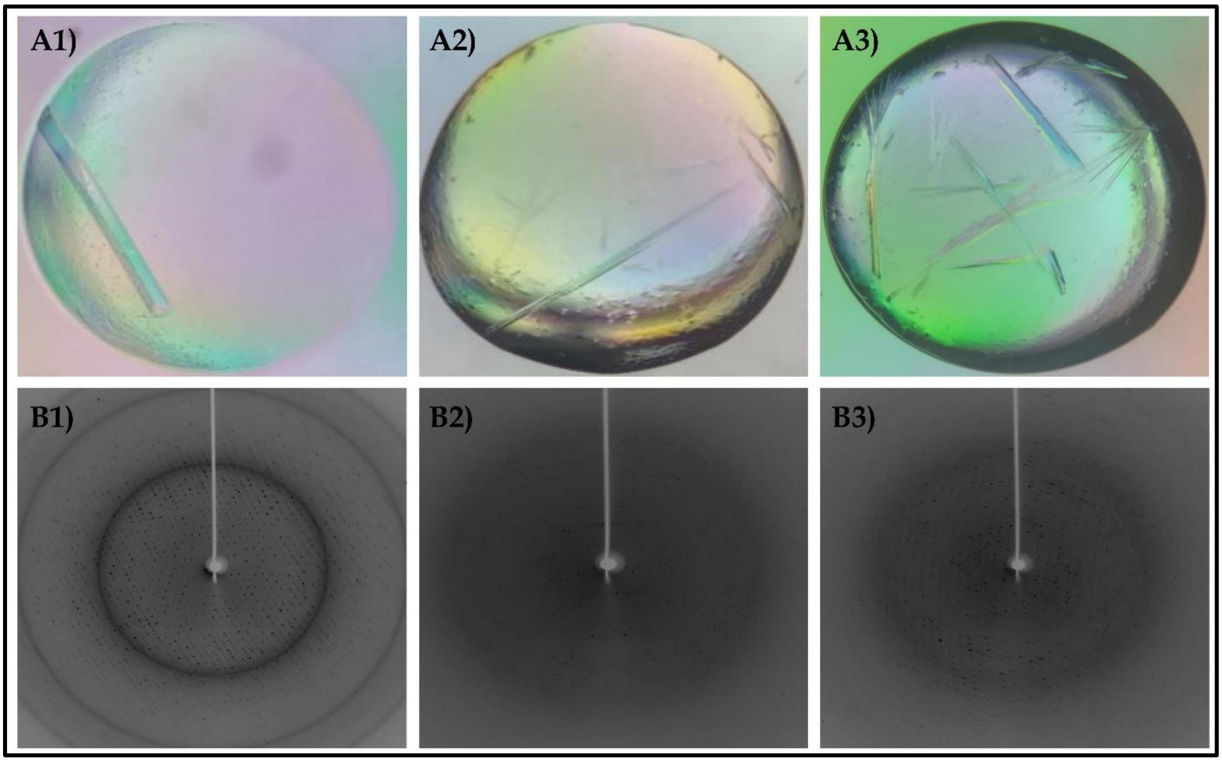


**Figure S1:** Crystallisation and diffractions. **A)** Crystals and **B)** their respective X-ray diffraction pattern. **A1)** and **B1)** PaIMPase-$Ca^{2+}$-2′AMP complex, **A2)** and **B2)** PaIMPase-$Mg^{2+}$ complex and PaIMPase-$Mg^{2+}$-Glycerotungstate complex, **A3)** and **B3)** PaIMPase-$Ca^{2+}$-MI-$PO_4^{3-}$ complex.

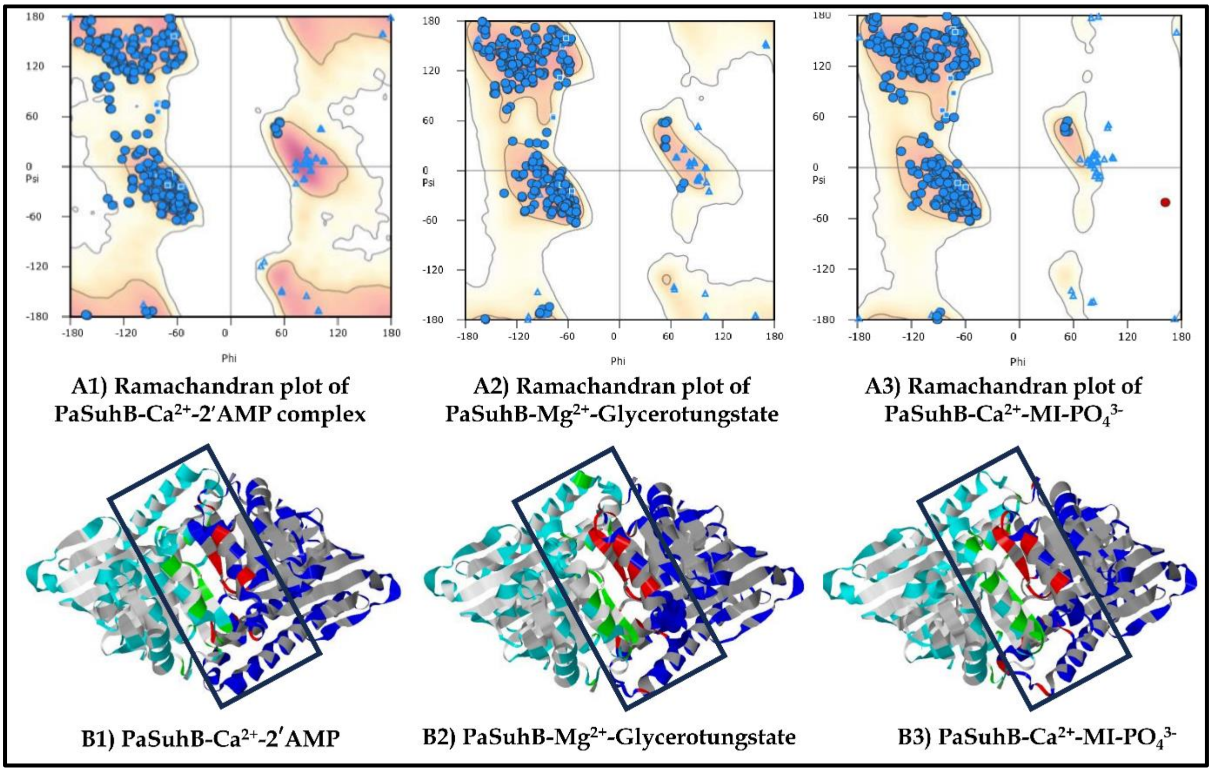


**Figure S2:** Model quality and dimer analysis. **A)** Ramachandran plot and **B)** their respective dimer interface area analysed by PISA server. **A1)** and **B1)** PaIMPase-$Ca^{2+}$-2′AMP complex, **A2)** and **B2)** PaIMPase-$Mg^{2+}$ complex and PaIMPase-$Mg^{2+}$-Glycerotungstate complex, **A3)** and **B3)** PaIMPase-$Ca^{2+}$-MI-$PO_4^{3-}$ complex.

**Table T1:** The interaction profile of the divalent cation at the active site of the PaSuhB structures in complex with substrate, transition state analogue and products.

| S. No. | Structure-complex | Metal, its position geometry and B factor (Å$^2$) | Coordinating atoms and respective distances |
|---|---|---|---|
| 1 | PaSuhB-$Ca^{2+}$-IPD (Chain A) | $Ca^{2+}$1 Octahedral, 22.72 | Asp86 (2.3 Å), Leu88 (2.3 Å), Glu67 (2.4 Å), IPD O8 (2.3 Å) and two waters (2.5 and 2.4 Å) |
| 2 | | $Ca^{2+}$2 Distorted octahedral, 34.90 | Asp86 (2.2 Å), Asp89 (1.8 Å), Asp216 (2.4 Å), IPD O1 (2.4 Å) and O8 (2.9 Å), one water (3.9 Å) |
| 3 | | $Ca^{2+}$3 Octahedral, 37.89 | Glu67 (2.4 Å), IPD O9 (2.4 Å), four waters (2.6, 2.5, 2.4 and 2.5 Å. |
| 4 | PaSuhB-$Ca^{2+}$-IPD (Chain B) | $Ca^{2+}$1 Octahedral, 24.15 | Asp86 (2.4 Å), Leu88 (2.4 Å), Glu67 (2.4 Å), IPD O9 (2.3 Å) and two waters (2.5 Å and 2.5 Å) |
| 5 | | $Ca^{2+}$2 Distorted octahedral, 42.12 | Asp86 (2.2 Å), Asp89 (2.4 Å), Asp216 (2.2 Å), IPD O1 (2.5 Å) and O9 (2.4 Å), one water (3 Å) |
| 6 | PaSuhB-$Ca^{2+}$-2′AMP (Chain A) | $Ca^{2+}$1 Octahedral, 42.44 | Asp86 (2.3 Å), Leu88 (2.3 Å), Glu67 (2.7 Å), 2AMP O1P (2.2 Å) and two waters (2.5 and 2.4 Å) |
| 7 | | $Ca^{2+}$2 Distorted octahedral, 49.92 | Asp86 (2.3 Å), Asp89 (2.4 Å), Asp216 (2.1 Å), 2AMP O1P (2.7 Å) and O2' (2.5 Å), one water (3.4 Å) |
| 8 | PaSuhB-$Ca^{2+}$-2′AMP (Chain B) | $Ca^{2+}$1 Octahedral, 37.63 | Asp86 (2.3 Å), Leu88 (2.5 Å), Glu67 (2.4 Å), 2′AMP O3P (2.2 Å) and two waters (2.4 and 2.5 Å) |
| 9 | | $Ca^{2+}$2 Distorted octahedral, 47.04 | Asp86 (2.2 Å), Asp89 (2.3 Å), Asp216 (2.2 Å), 2′AMP O2′ (2.5 Å) and O3P (2.6 Å), one water (3.2 Å) |
| 10 | PaSuhB-$Mg^{2+}$-GTG (Chain A) | $Mg^{2+}$1 Tetrahedral, 45.66 | Asp86 (1.9 Å), Leu88 (2.0 Å), Glu67 (1.9 Å), GTG O05 (2.4 Å) and O03 (2.7 Å) |
| 11 | | $Mg^{2+}$2 Tetrahedral, 44.30 | Asp86 (2.2 Å), Asp89 (2.1 Å), Asp216 (1.9 Å), GTG O05 (1.8 Å) and one water (3.1 Å) |
| 12 | PaSuhB-$Mg^{2+}$-GTG (Chain B) | $Mg^{2+}$1 Distorted octahedral, 47.44 | Asp86 (2.0 Å), Leu88 (2.5 Å), Glu67 (2.0 Å), GTG O05 (2.6 Å) and O03 (2.9 Å) and one water (2.2 Å) |
| 13 | | $Mg^{2+}$2 Tetrahedral, 43.92 | Asp86 (2.2 Å), Asp89 (2.1 Å), Asp216 (2.5 Å), and GTG O05 (1.9 Å) |
| 14 | PaSuhB-$Mg^{2+}$-MI-$PO_4^{3-}$ (Chain A) | $Ca^{2+}$2 Distorted octahedral, 50.99 | Asp86 (2.2 Å), Asp89 (2.7 Å), Asp216 (2.2 Å), and O2 of MI (2.9 Å), and two waters (2.4 and 2.7 Å) |
| 15 | PaSuhB-$Mg^{2+}$-MI-$PO_4^{3-}$ (Chain B) | $Ca^{2+}$2 Distorted octahedral, 46.22 | Asp86 (2.4 Å), Asp89 (2.5 Å), Asp216 (2.2 Å), and O2 of $PO_4^{3-}$ (2.2 Å), and two waters (2.4 and 2.8 Å) |

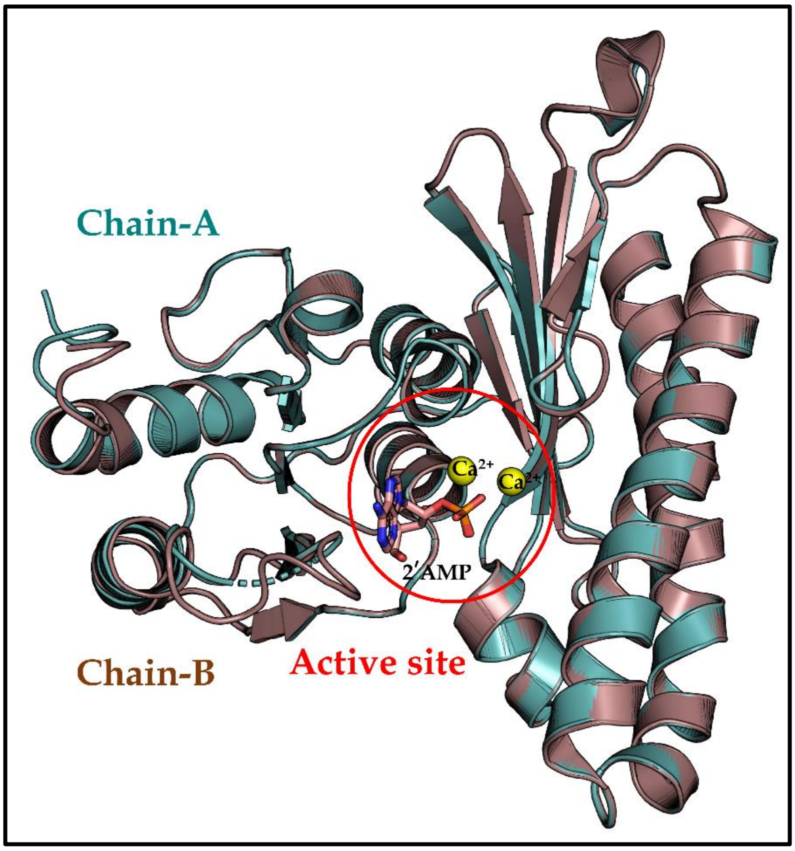


**Figure S3:** Overlapped view of two monomers of PaIMPase-$Ca^{2+}$-2′AMP complex crystal structure

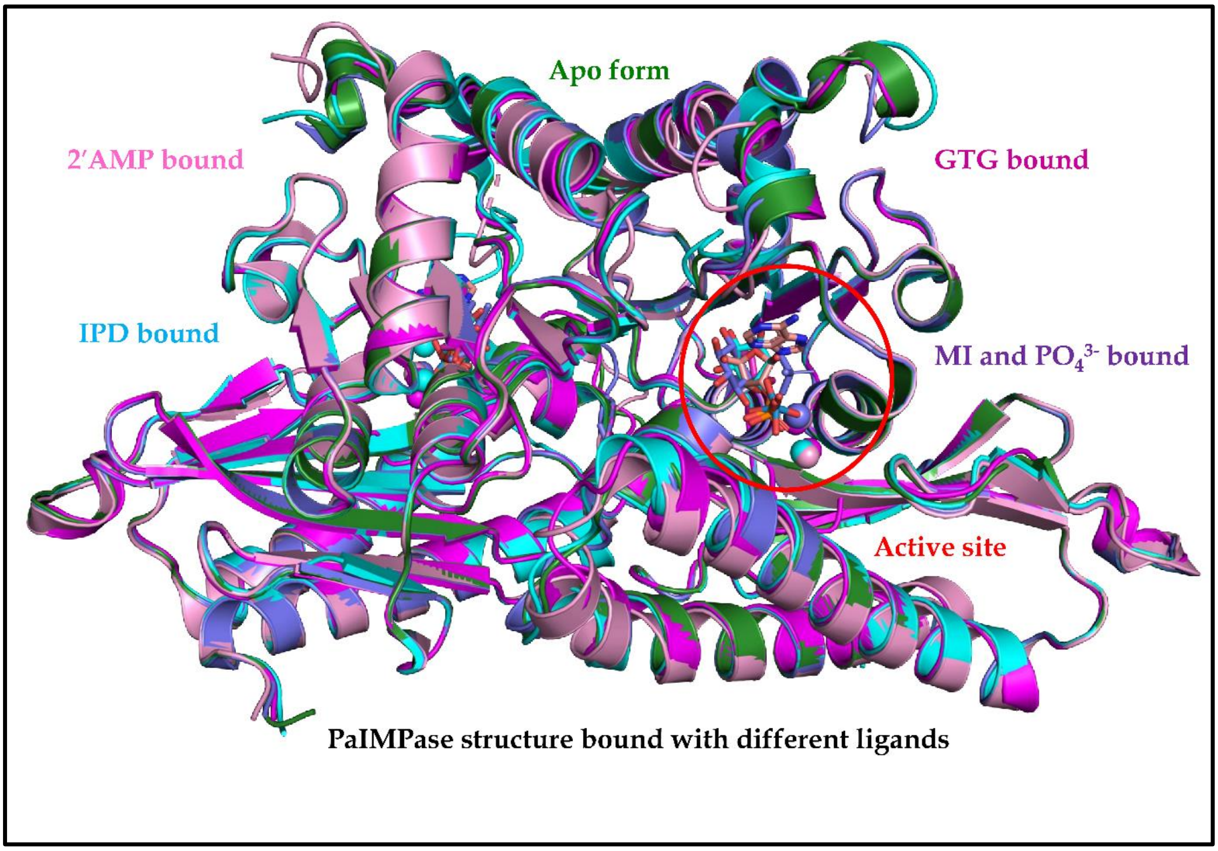


**Figure S4:** Overlapped view of the dimeric structures, including apo, 2′AMP, IPD, GTG, and products MI and $PO_4^{3-}$-bound, representing the conserved dimeric structures with lesser Cα RMSD values.

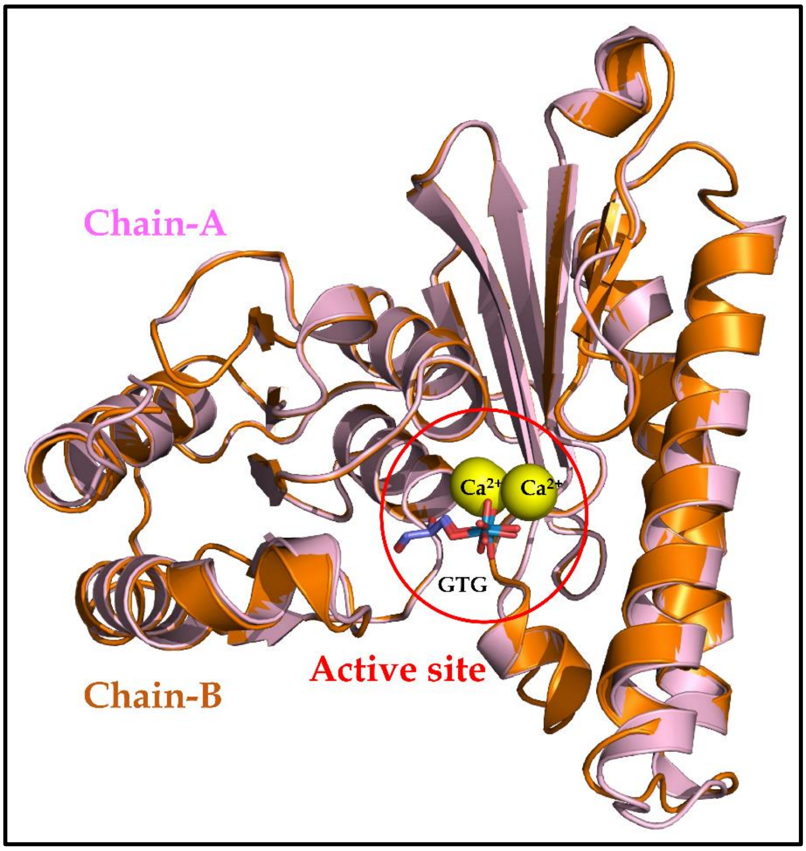


**Figure S5:** Overlapped view of two monomers of PaIMPase-$Mg^{2+}$-GTG complex crystal structure

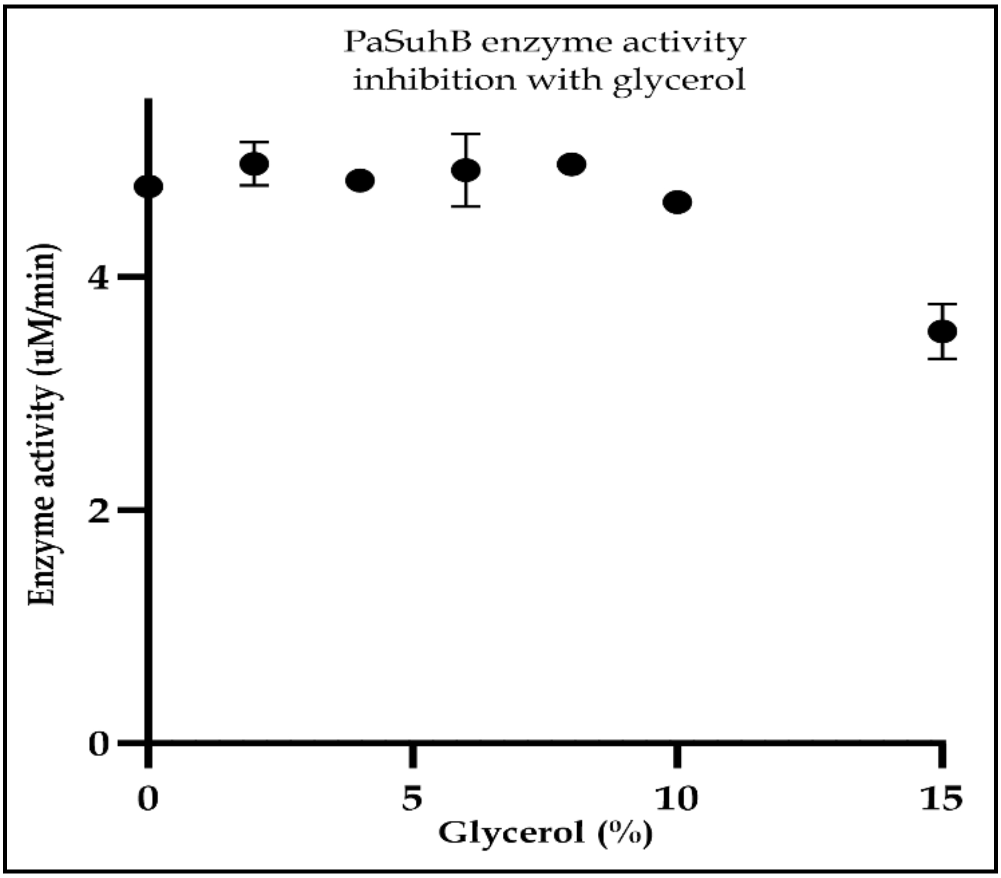


**Figure S6:** Glycerol-based PaSuhB enzyme activity inhibition

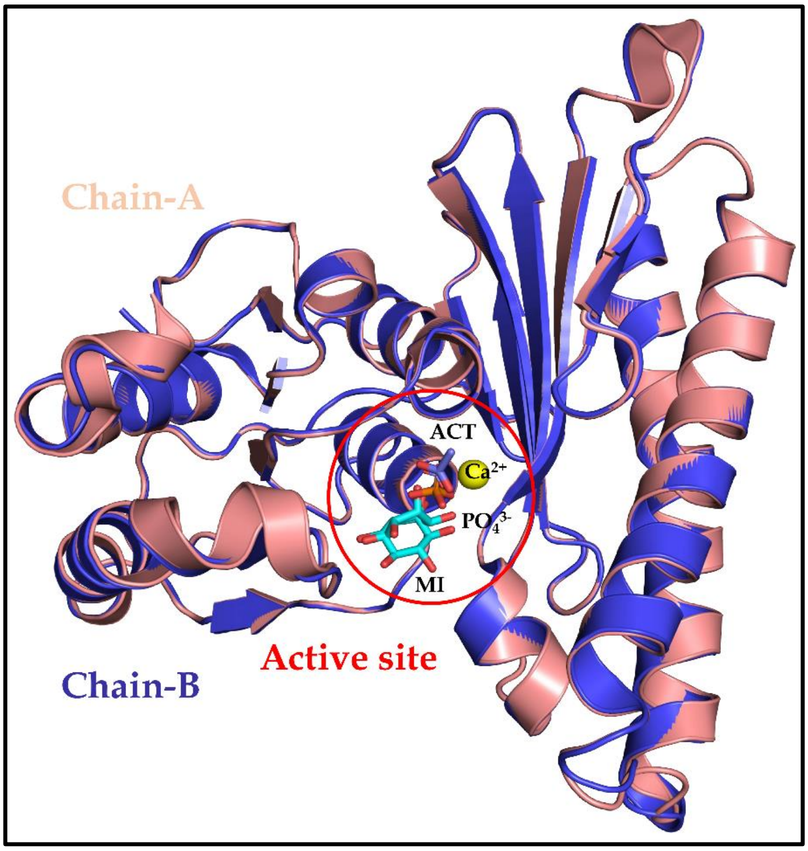


**Figure S7:** Overlapped view of two monomers of PaIMPase-$Ca^{2+}$-MI-$PO_4^{3-}$ complex crystal structure

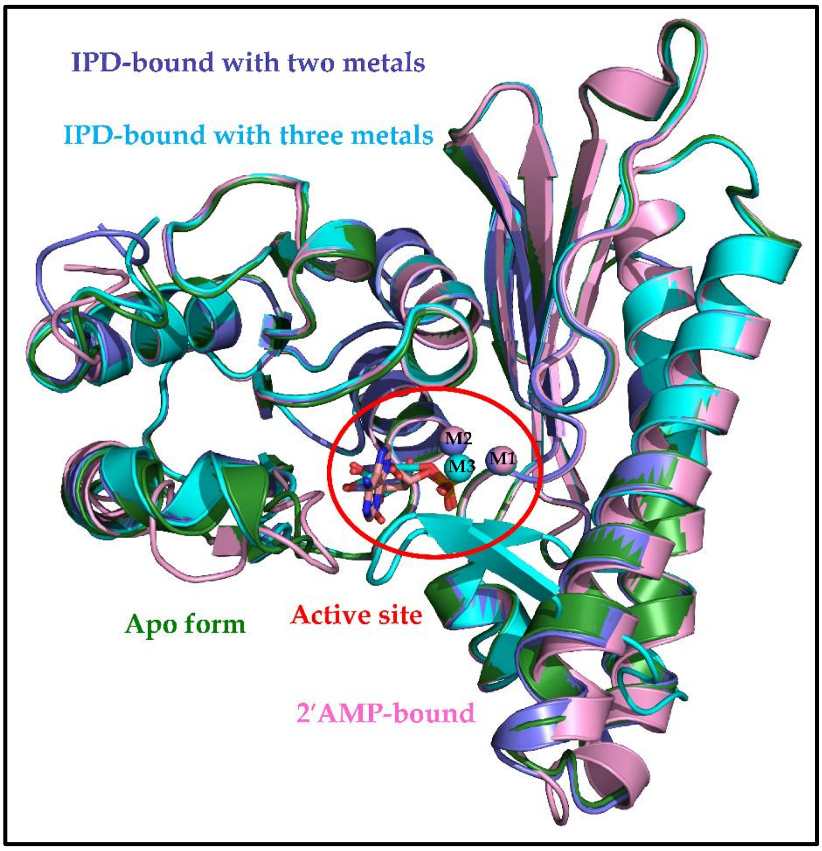


**Figure S8:** Overlapped view of the monomers of the apo and substrate-bound crystal structures showing overall 3D fold conservation.

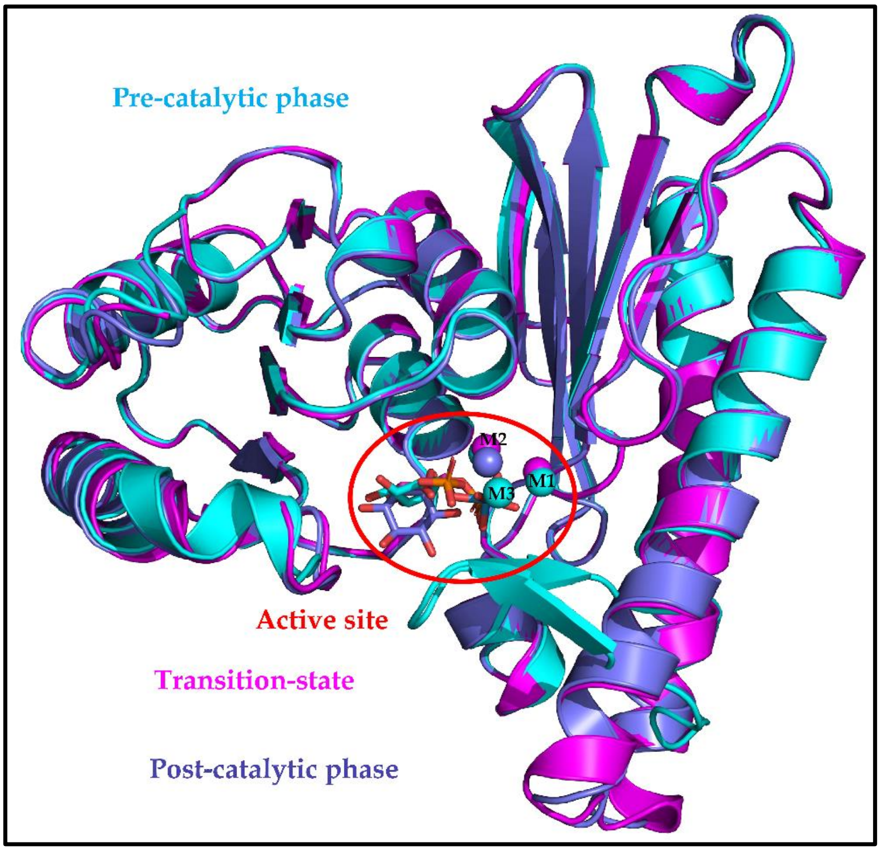


**Figure S9:** Overlapped view of the monomers representing precatalytic, transition-state and post-catalytic events showing overall 3D fold conservation with minimal Cα RMSD differences.

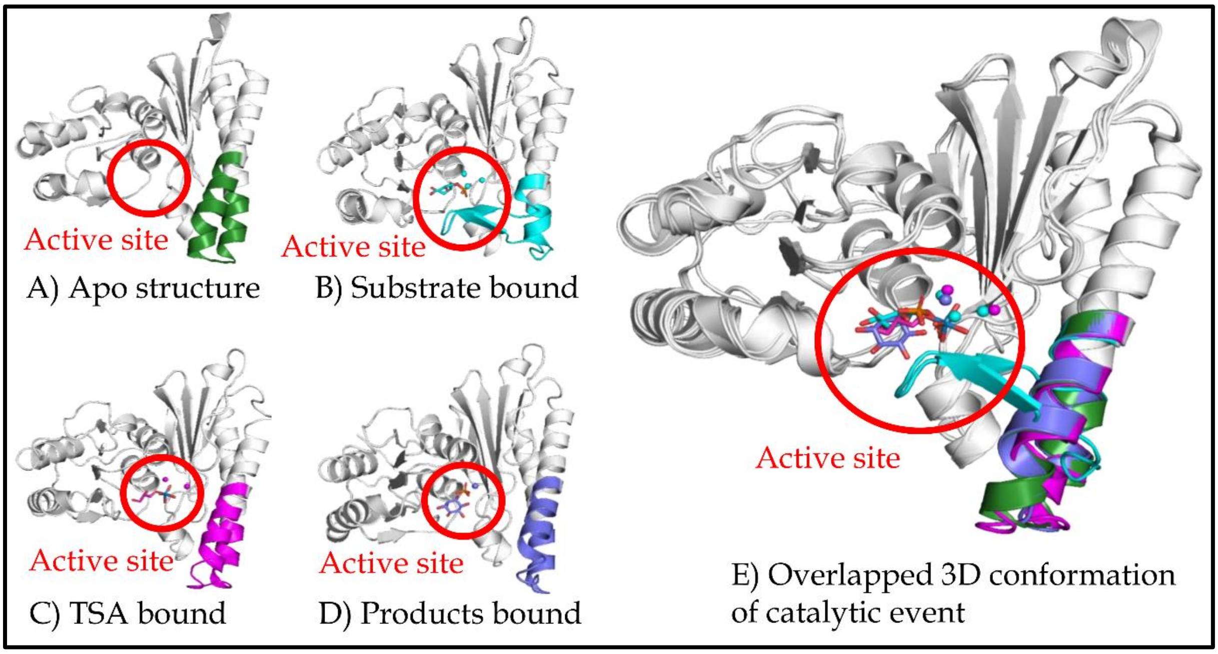


**Figure S10:** 3D orientation of active site mobile loop at the different conformational stages of PaIMPase enzymatic catalysis. **A)** The apo-PaIMPase structure showing the empty active site, **B)** Substrate (IPD) bound PaIMPase structure showing the precatalytic phase, **C)** Transition state bound PaIMPase structure showing the intermediate state of the catalysis, **D)** Products bound PaIMPase structure showing the post-catalytic phase of the reaction, **E)** Overlapped 3D conformations of PaIMPase monomers representing different catalytic events at the active site enzyme.

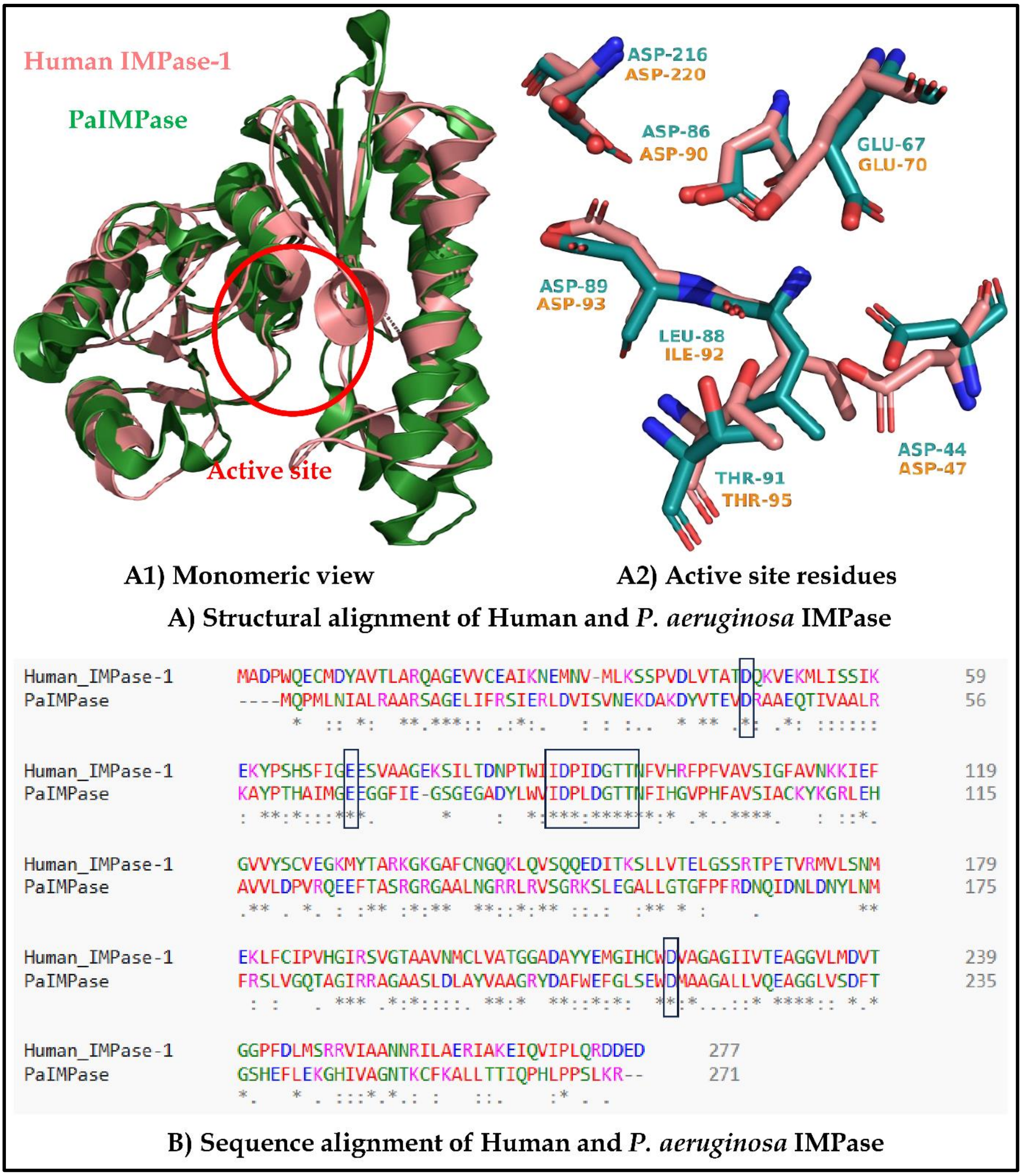


**Figure S11:** Structure and sequence comparison of human and *P. aeruginosa* IMPase, **A)** Structural alignment showing **A1**) overall 3D fold conservation through monomeric structure and **A2)** Conserved residues at the active site, **B)** Sequence alignment showing the conserved active site amino acid residues.

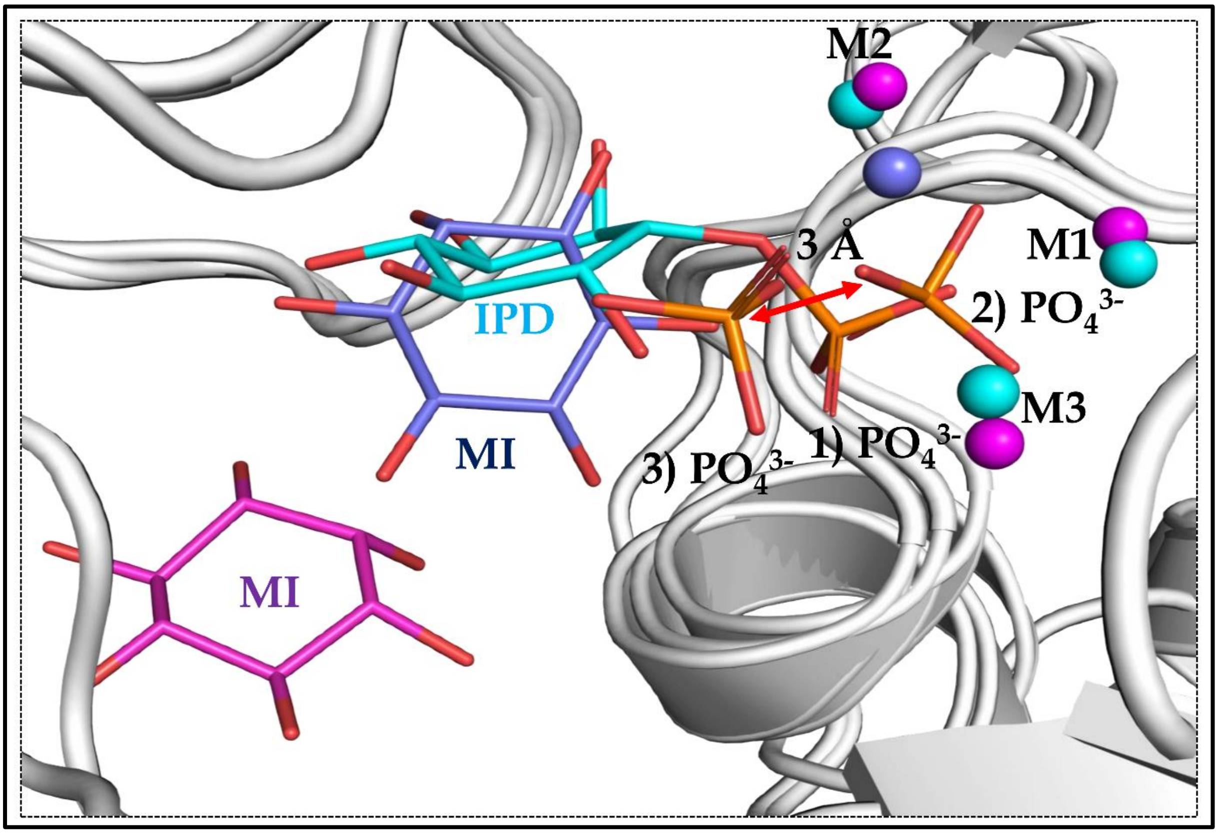


**Figure S12:** Comparative 3D positioning of the $PO_4^{3-}$ from substrate (PDB: 8WDQ), immediate product (PDB: 1G0I), and collapsed product (PDB: 9XCK) bound crystal structures. M1, M2, and M3 represent the position of three metal ions. Colour coding is used for the structures, such as substrate (Cyan), immediate product (Magenta), and collapsed product (Blue). $PO_4^{3-}$ is numbered based on the sequential events happening at the active site; the substrate, immediate product, and collapsed product states of $PO_4^{3-}$ are given numbers 1, 2, and 3, respectively.
